# A new behavioral and neural marker of social vision

**DOI:** 10.1101/2021.06.09.447707

**Authors:** Etienne Abassi, Liuba Papeo

**Author notes:** Correspondence to: E.A., L.P. **Competing interests:** All authors declare that they have no financial or non-financial competing interests.

## Abstract

Research on face perception has revealed highly specialized visual mechanisms such as configural processing, and provided markers of interindividual differences –including disease risks and alterations– in visuoperceptual abilities that traffic in social cognition. Is face perception unique in degree or kind of mechanisms, and in its relevance for social cognition? Combining functional MRI and behavioral methods, we address the processing of an uncharted class of socially relevant stimuli: minimal social scenes involving configurations of two bodies spatially close and face-to-face as if interacting (hereafter, facing dyads). We report category-specific activity for facing (*vs*. non-facing) two-body configurations in selective areas of the visual cortex. That activity shows face-like signatures of configural processing –i.e., stronger response, and greater susceptibility to stimulus inversion for facing (*vs*. non-facing) dyads–, and is predicted by performance-based measures of body-dyad perception (i.e., accuracy in a fast visual categorization task). Moreover, individual performance in body-dyad perception is reliable, stable-over-time and correlated with the individual social sensitivity, coarsely captured by the Autism-Spectrum Quotient. Further analyses clarify the relationship between single-body and body-dyad perception. We propose that facing dyads are processed through highly specialized mechanisms (and brain areas), analogously to other biologically/socially relevant stimuli such as faces. Like face perception, facing-dyad perception can reveal basic visual processes that lay the foundations for understanding others, their relationships and interactions.

**Significance statement:** With its specialization to faces and biological motion, vision houses the foundations of human social ability. Using a multimodal approach (meta-analysis, fMRI, visual-perception tasks and self-administered survey), we introduce a new class of visual stimuli –minimal social scenes with two face-to-face bodies–, whose processing highlights new behavioral and neural markers of visuoperceptual abilities that traffic in social cognition. Behavioral and neural effects of body-dyad perception reveal the recruitment of specialized configural processing, previously described for face perception. Furthermore, individual performance in body-dyad perception is stable over time, and predicts an individual’s social sensitivity, measured in terms of autism-spectrum traits. Thus, body-dyad perception reveals uncharted aspects of visual functioning and specialization, which may critically contribute to human social life.

## Introduction

Few functions of the human brain have been explored as much and deeply as the visual processing of faces. And few visual functions appear as vital for social life as face processing. Face processing underlies detection, recognition and identification of conspecifics, and encoding of visual information that provides the basis for attributing states, traits, goals and intentions to others, and regulating one’s own behavior in social contexts (1, 2). Thus, it is not surprising that highly efficient and specialized mechanisms exist for face perception, in the human visual system.

One of the most robust markers for specialized face processing mechanisms is the *face inversion effect* (FIE), a disproportionate cost in processing a face inverted upside-down relative to a face in the canonical upright orientation (3). The FIE has revealed a visual mechanism, so-called configural, that treats the face not as a set of parts (all present in both upright and inverted stimuli), but as a perceptual unit with a specific configuration (i.e., eyes above mouth), which is disrupted by inversion, yielding catastrophic effects on recognition (4, 5).

Besides revealing important aspects of visual functioning, the sensitivity to face inversion is reliable and stable over time at the intraindividual level, and can highlight interindividual differences that capture the normal variability in the general population, as well as disease risks and alterations in visual perception (6, 7) and in socially relevant cognitive abilities (8–10). Reduced sensitivity to face inversion –and atypical face processing in general– is seen in developmental disorders that affect social cognition such as the autism spectrum disorder (ASD) (8, 11–16), although with high variability within the ASD population and no consensus on the neurofunctional basis of the alteration (17–21).

*Is face perception unique in degree or kind of cognitive and neural mechanisms, and in its relevance for social cognition?* While faces remain the best understood case of visual specialization, other visual stimuli that also support social cognitive tasks, such as bodies and biological motion (2), may recruit analogous mechanisms (22, 23). Moreover, recent research indicates an uncharted class of stimuli, whose visual processing might be analogous to faces. Members of this class are two-body shapes, that is, configurations of two bodies spatially close and face-to-face, as is (proto)typical of social interaction (24, 25). Much of what we learn about social life, and how we regulate our social behavior *here and now*, depends on the observation of people interacting with people (26, 27). Thus, the visual specialization for face-to-face body dyads (hereafter, facing dyads) might respond to the need for efficient detection and recognition of social interaction in the cluttered and crowded visual world.

It has been suggested that visual processing of facing dyads share with face processing a particular sensitivity to inversion: In fast visual categorization (i.e., discrimination of bodies from other objects), subjects’ performance drops dramatically for inverted facing dyads, relative to the same dyads presented upright (24, 28). The cost of inversion for facing dyads is significantly larger than the cost of inversion for identical bodies in a non-facing configuration (e.g., back-to-back bodies), an effect known as the two-body inversion effect (2BIE).

*What is the two-body inversion effect?* The present study asked whether, like the FIE, the 2BIE is *a)* an index of visual specialization mediated by configural processing, and *b)* a reliable measure of the individual visuoperceptual functions that traffic in social perception and cognition.

To address the first question, we tested whether activity in body-specific visual areas of the occipitotemporal cortex showed signatures of configural processing during perception of facing dyads. A neural signature of configural face processing is greater activity for faces relative to scrambled or scattered faces (29). Thus, if facing dyads recruit configural processing, they should evoke greater activity in body-specific visual areas, relative to their scattered counterparts, i.e., stimuli featuring the same parts (bodies) in a non-facing configuration. Moreover, the 2BIE captured in the subjects’ behavior should predict the activity of body-specific visual areas in response to upright *vs*. inverted dyads, just like the FIE predicts, across subjects, the response to upright *vs*. inverted faces in the face-specific visual cortex (30). To test these predictions, we recorded neural activity using functional MRI (fMRI), while healthy subjects viewed upright or inverted facing and non-facing dyads. On a different day, the same subjects performed a visual categorization task to measure the 2BIE (24). We tested the visual specificity of the whole-dyad configuration considering the difference in neural activity for facing *vs*. non-facing dyads, and the correlation between the behavioral 2BIE and the effect of inversion on the neural response to dyads (the neural 2BIE) in individual subjects.

To address the second question, subjects performed the visual categorization task a second time after weeks, so that we could estimate the stability of the 2BIE over time. They also completed the Autism-Spectrum Quotient Test (AQ), a measure of the autism-spectrum traits, sensitive to interindividual differences in the general population, with respect to socially relevant cognitive abilities (31, 32). Significant correlations between the two measures of the 2BIE (i.e., test-retest reliability) and between the 2BIE and the AQ, at the individual level, will provide first indication that visual perception of social scenes can be a reliable measure of the individual’s visuoperceptual functions that contribute to lay the foundations for understanding others relationships and interactions.

## Results

### Processing of body dyads

#### The behavioral 2BIE

To measure the behavioral 2BIE (i.e., the larger cost of inversion on the recognition of facing dyads, relative to non-facing dyads; 24), 22 subjects of the 29 included in the fMRI study, took part in two identical sessions of the same visual categorization task. During this task, they saw images of single bodies, facing or non-facing dyads, and of single chairs, and facing or non-facing pairs of chairs. All images were presented upright or inverted upside-down, for a short time (30 ms), and then masked. For each trial, subjects had to report whether they had seen bodies or chairs, regardless of the number of items (one/two), positioning (facing/non-facing) or orientation (upright/inverted). The second session took place on average 20 days after the first. Accuracy data averaged across the two sessions of the task were analyzed in a 2 Category (body, chair) x 3 Stimulus (facing, non-facing, single stimuli) x 2 orientation (upright, inverted) repeated-measures ANOVA. Results showed a trend for a main effect of Category [*F*(1,21) = 3.82, *p* = 0.064, *η*_p_^2^ = 0.15], significant effects of Stimulus [*F*(2,42) = 7.67, *p* = 0.001, *η*_p_^2^ = 0.27] and Orientation [*F*(1,21) = 25.63, *p* < 0.001, *η*_p_^2^ = 0.55], and significant interactions between Category and Stimulus [*F*(2,42) = 8.45, *p* < 0.001, *η*_p_^2^ = 0.29], Category and Orientation [*F*(1,21) = 14.1, *p* = 0.001, *η*_p_^2^ = 0.40], and Stimulus and Orientation [*F*(2,42) = 6.27, *p* = 0.004, *η*_p_^2^ = 0.23]. Main effects and interactions were qualified by a significant three-way interaction between Category, Stimulus and Orientation [*F*(2,42) = 9.13, *p* = 0.001, *η*_p_^2^ = 0.30]. Two separate ANOVAs with body-trials only and chair-trials only, showed that the magnitude of the inversion effect was affected by the type of Stimulus (single, facing, non-facing), for body-stimuli (Effect of Stimulus: [*F*(2,42) = 9.51, *p* < 0.001, *η*_p_^2^ = 0.31]; Effect of Orientation: [*F*(1,21) = 21.25, *p* < 0.001, *η*_p_^2^ = 0.50], Interaction: [*F*(2,42) = 10.78, *p* < 0.001, *η*_p_^2^ = 0.34]), but not for chair-stimuli (Effect of Stimulus: [*F*(2,42) = 3.19, *p* = 0.051, *η*_p_^2^ = 0.13]; Effect of Orientation: [*F*(1,21) = 5.27, *p* = 0.032, *η*_p_^2^ = 0.20], Interaction: [*F*(2,42) = 1.61, *p* = 0.212, *η*_p_^2^ = 0.07]; see *Supplemental Information* for details on the performance with chairs).

Considering more closely the performance with body-stimuli, pairwise *t*-tests showed that, while the inversion effect (i.e., the difference between upright *vs*. inverted stimuli) was significant for all three stimulus conditions [facing dyads: *t*(21) = 4.88, *p* < 0.001; non-facing dyads: *t*(21) = 4.28, *p* < 0.001; single bodies: *t*(21) = 4.10, *p* = 0.001], it was larger for facing dyads, relative to non-facing dyads, *t*(21) = 3.53, *p* = 0.002, and single bodies, *t*(21) = 3.76, *p* = 0.001, and comparable in the last two conditions, *t*(21) = 1.21, *p* = 0.240 (Fig. 1a). Separate analyses for data collected in the first and in the second session of the task gave consistent results (*Supplemental Information 1,2* and Fig. S1a). Consistent with previous reports (24, 33, 34), accuracy proved more sensitive than reactions times (RTs) to the effects of inversion in the current task. RT analyses gave no effects or conformed to the pattern of accuracy results (*Supplemental Information 1, 2, 4;* Fig. S1b-c).

**Figure 1.**
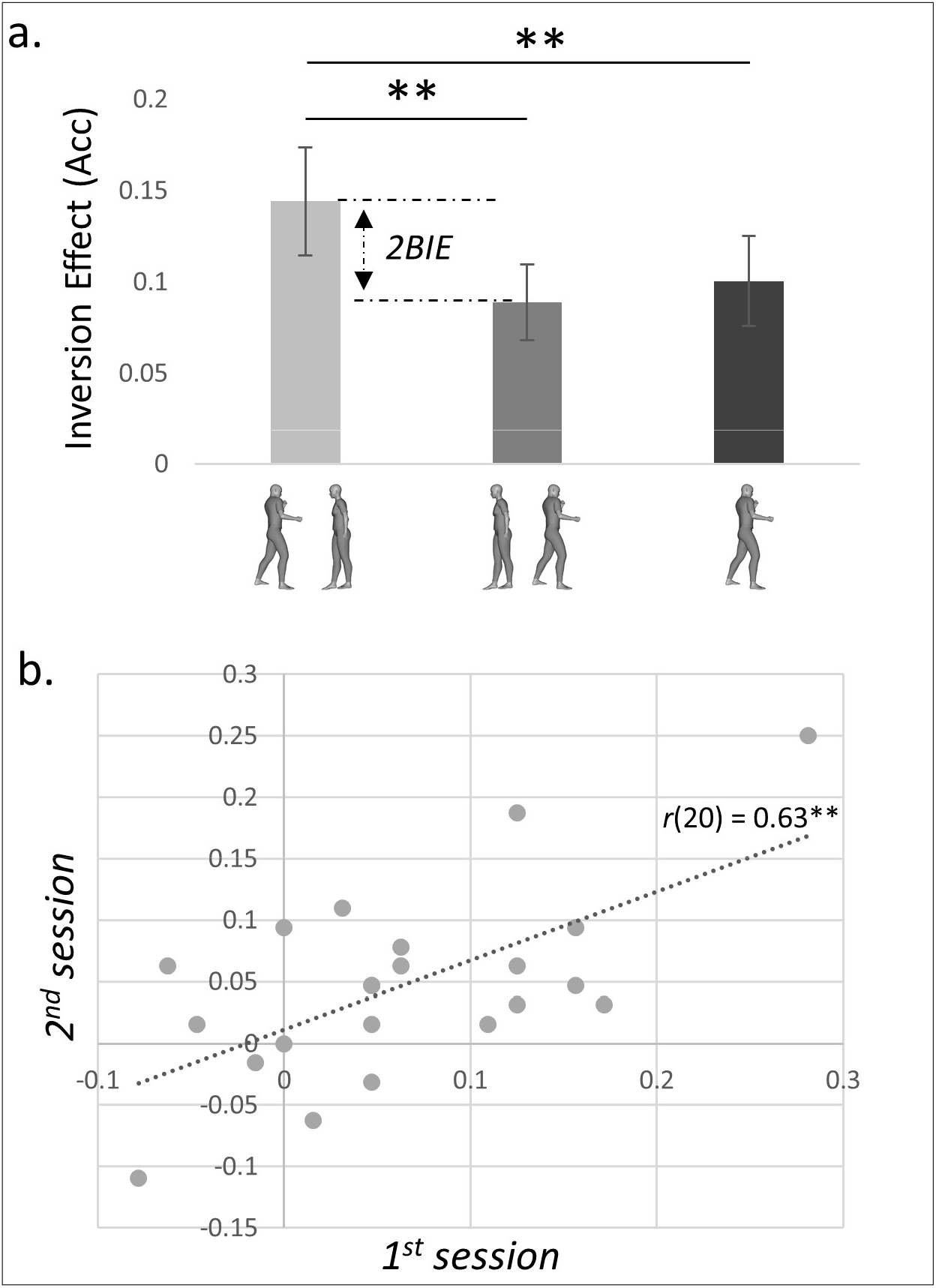
Behavioral 2BIE and within-subject correlation between the two measures of the 2BIE. **a**. Inversion effect (proportion of correct responses for upright minus inverted trials) averaged over the two behavioral sessions for facing dyads, non-facing dyads and single bodies. Error bars denote the within-subjects normalized standard error of the mean; \*\**p* ≤ 0.01. **b**. Pearson correlation between the size of the 2BIE for the same subjects, measured in the first and second session of the visual categorization task. \*\**p* ≤ 0.01.

At the group level, there was no difference in the magnitude of the 2BIE between the first and the second session [*t*(21) = 1.13, *p* > 0.250, two-tail], although accuracy was overall higher in the second session [*t*(21) = 2.96, *p* = 0.008, two-tail], most likely as an effect of learning. The test-retest reliability of the 2BIE was computed considering, for each subject, the two measures of the 2BIE [(Upright-inverted)_facing_-(Upright-inverted)_non-facing_], measured in the behavioral session 1 and 2. We found a significant within-subjects Pearson correlation [*r*(20) = 0.63, p = 0.002], which was significantly higher than any random between-subjects correlation [*p* = 0.001] (Fig. 1b). This result defines the 2BIE as a reliable, stable over time, measure of the individual visual processing of body dyads.

Finally, we compared, qualitatively, the size of the inversion effects for facing and non-facing dyads, as measured in the current study and in three previous studies (24, 33, 34), to the size of the inversion effects for single faces, bodies and objects, as measured in various previous studies reviewed in Rezlescu et al. (35) (The full list of studies is provided as in Table S2 of Supplementary Information). To this end, the scaled inversion effects (%) were computed for all the studies as [(raw inversion effect) / (1-chance)], and visualized in a single plot. Average scaled inversion effects are (in descending order): 28.4 ±8.55 SD for facing bodies (4 studies), 22.78 ±13.78 for faces (55 studies), 18.5 ±7.07 for non-facing bodies (4 studies), 17.71 ±7.74 for single bodies (9 studies), 4.76 ±6.63 for other objects (56 studies). Thus, as shown in Figure 2, the inversion effect for facing body dyads appears systematically larger than the inversion effect for non-facing dyads and closer to the inversion effect for faces than all other categories (single bodies and other objects).

**Figure 2.**
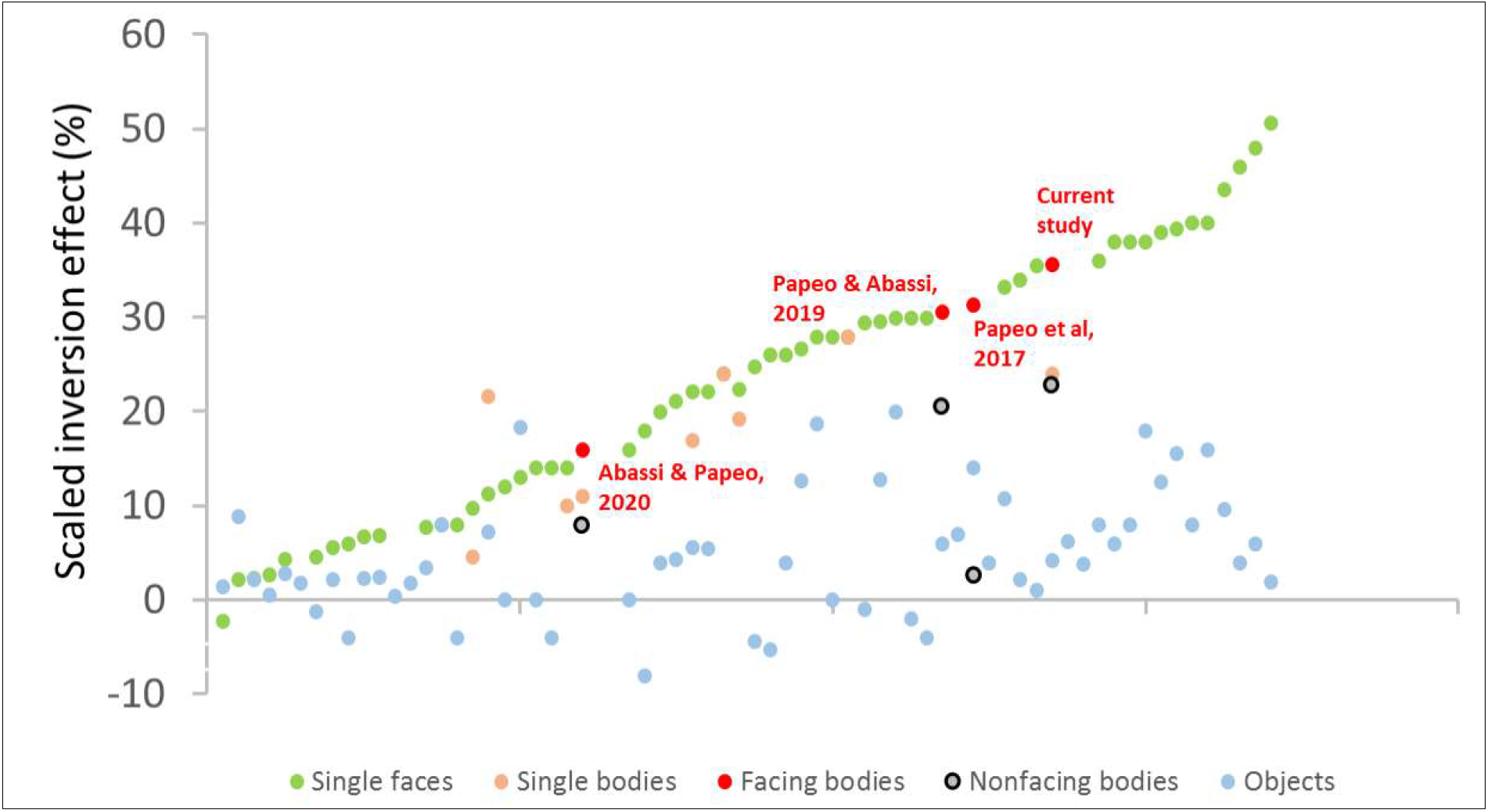
Scaled inversion effect (%) for facing dyads compared to the effect for non-facing dyads, single faces, single bodies and other objects. Each point represents the scaled inversion effect measured in one of the studies reviewed in Rezlescu et al. (2016), to which we added values of the scaled inversion effect for facing and non-facing body-dyads, as measured in three previous studies and in the current one. Studies are sorted on the X axis according to the size of the scaled inversion effect for each category of stimuli in each study. written in red on the figure correspond to the closest red dot and denote the size of inversion effect for facing dyads.

#### Neural signatures of visual configural processing

A neural signature of configural face processing is the increased response to faces (or bodies) *vs*. scattered stimuli in category-specific visual areas of the occipitotemporal cortex (29, 36). The current results showed an analogous effect during processing of body dyads, where non-facing dyads were taken as the scattered counterpart of facing-dyad configurations. Converging evidence for increased response to facing *vs*. non-facing dyads was obtained with whole-brain and region of interest (ROI) analyses.

The first analysis was based on data collected during the main fMRI experiment, in which 29 subjects in the fMRI scanner viewed upright or inverted images of facing or non-facing body dyads, and, in independent runs, upright or inverted images of single bodies or objects. The whole-brain contrast [upright facing > non-facing dyads] showed an effect in a cluster centered in the left posterior middle temporal gyrus, extending into the lateral occipital and overlapping with the body-specific extrastriate body area (EBA) [MNI peak coordinates: -50 -72 12; peak *z*-value = 3.96; peak *p*-value < 0.001; cluster size = 336; *p* = 0.013, FWE-corrected; see Fig. 3b]. For the contrast [inverted facing > non-facing dyads], no clusters survived the FWE correction.

**Figure 3.**
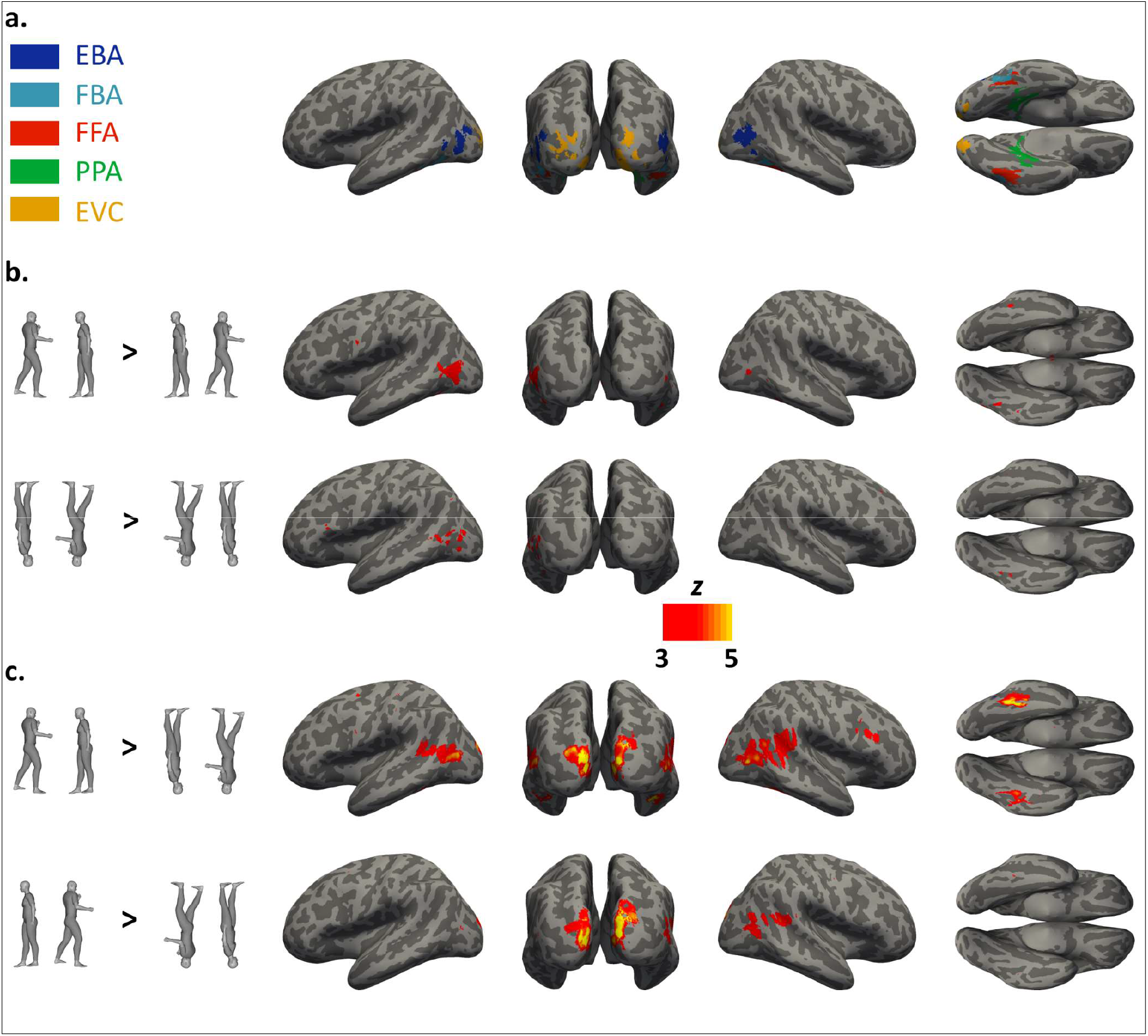
Definition of Regions of Interest (ROIs) and results of whole-brain contrasts addressing the effects of configural processing in body-dyad perception. **a**. ROI location on an inflated brain, based on fMRI data registered during the functional localizer task: EBA (extrastriate body area), FBA (fusiform body area), FFA (fusiform face area), PPA (parahippocampal place area) and EVC (early visual cortex). Group-level clusters are shown for illustration purposes. **b**. Group random-effect maps (*n* = 29) for the contrast upright facing > non-facing dyads (upper row) and inverted facing > non-facing dyads (lower row). **c**. Group random-effect maps (*n* = 29) for the contrast upright > inverted facing dyads (upper row) and upright > inverted non-facing dyads (lower row). The color bar indicates uncorrected *z* values.

The above effect of body positioning in the occipitotemporal cortex was confirmed by an ROI analysis. In particular, using an independent functional localizer task (Fig. 3a) during fMRI, we identified, for each subject, two areas of the visual occipototemporal cortex showing the highest response to body stimuli, the EBA and the fusiform body area (FBA). In addition, we localized the individual face-selective fusiform face area (FFA). The FFA was localized not only because faces were part of our stimuli, but also because this area has consistently been implicated in configural processing of faces as well as bodies (29, 30). Finally, to control for the specificity of effects in the above ROIs, we considered two additional ROIs, in the place-specific parahippocampal place area (PPA), and in the early visual cortex (EVC). From each ROI, we extracted the activity (percentage of signal change, PSC) for upright and inverted, facing and non-facing body dyads. A 5 ROIs x 2 Stimulus (facing, non-facing) x 2 Orientation (upright, inverted) ANOVA showed significant effects of ROI [*F*(4,112) = 69.89, *p* < 0.001, *η*_p_^2^ = 0.71], Stimulus [*F*(1,28) = 4.85, *p* = 0.036, *η*_p_^2^ = 0.15] and Orientation [*F*(1,28) = 23.92, *p* < 0.001, *η*_p_^2^ = 0.46], and significant interactions between ROI and Stimulus [*F*(4,112) = 12.74, *p* < 0.001, *η*_p_^2^ = 0.31], and ROI and Orientation [*F*(4,112) = 10.65, *p* < 0.001, *η*_p_^2^ = 0.28], but not between Stimulus and Orientation [*F*(1,28) = 1.09, *p* > 0.250, *η*_p_^2^ = 0.03]. All significant effects and interactions were qualified by a significant three-way interaction [*F*(4,112) = 2.69, *p* = 0.035, *η*_p_^2^ = 0.09], showing that the difference in the magnitude of the inversion effects on neural activity for facing and non-facing dyads varied across ROIs.

Next, we used pairwise *t*-tests to assess, in each ROI, the two signatures of configural processing targeted here: 1) larger response to facing than to non-facing (upright) dyads, and 2) larger difference between upright and inverted stimuli for facing, than for non-facing dyads (or the neural 2BIE). We found stronger response to upright facing than non-facing dyads in the FFA [*t*(28) = 3.41, *p* < 0.001], EBA [*t*(28) = 3.52, *p* < 0.001] and FBA [*t*(28) = 2.51, *p* = 0.009], but not in the PPA [*t*(28) = 0.09, *p* > 0.250] and EVC [*t*(28) = 0.08, *p* > 0.250] (Fig. 4a). Moreover, we found a larger inversion effect for facing than for non-facing dyads in the FFA [*t*(28) = 2.36, *p* = 0.013], and a trend in the same direction in the EBA [*t*(28) = 1.49, *p* = 0.073]. No other ROI showed a similar effect or trend [FBA: *t*(28) = 0.88, *p* = 0.192; PPA: *t*(28) = 0.67, *p* > 0.250; EVC: *t*(28) = 0.74; p = 0.234] (Fig. 4b) (for whole-brain contrasts addressing the effects of inversion, see Fig. 3c; Table S1).

**Figure 4.**
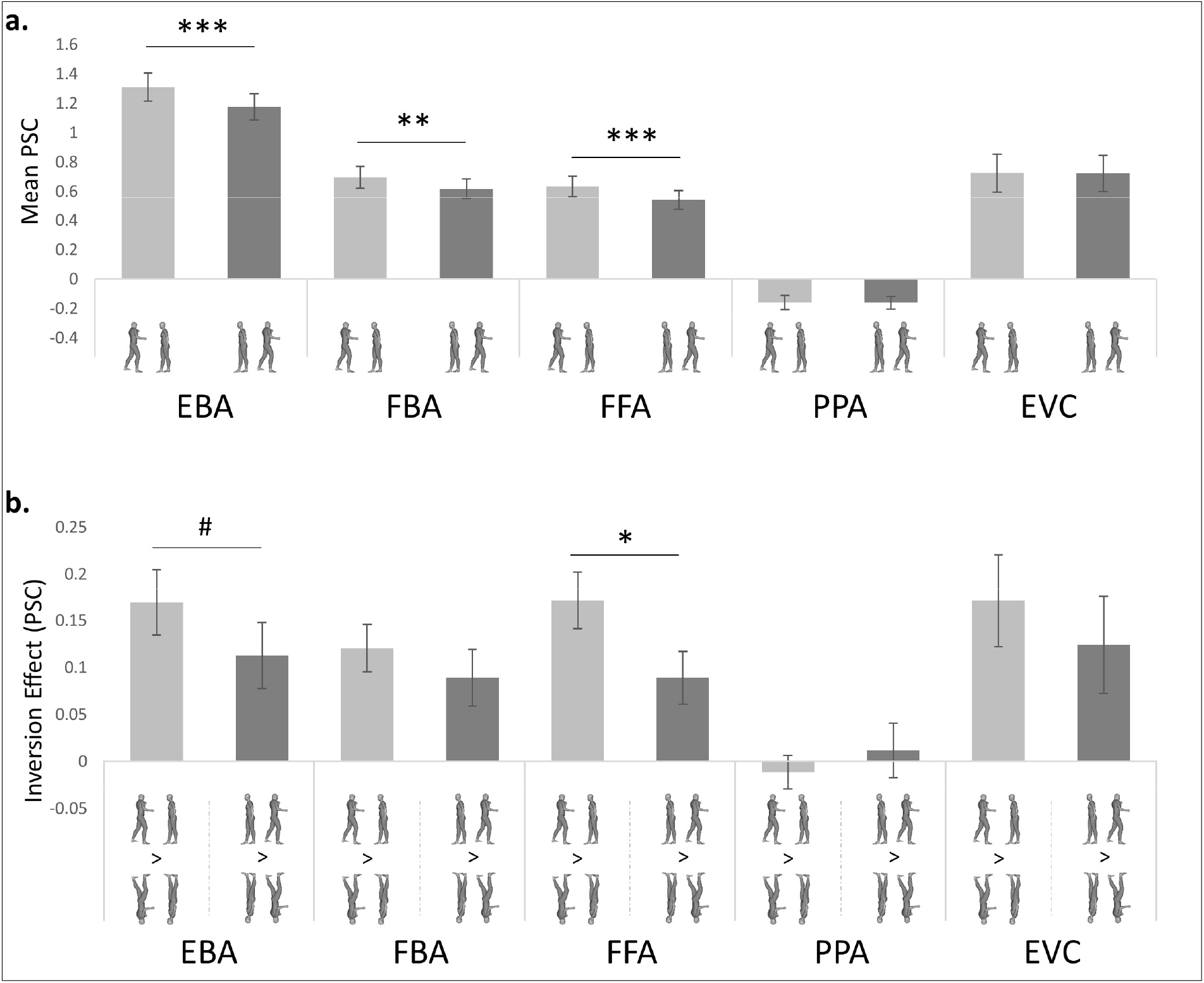
Neural markers of configural processing for body dyads in ROIs. Analyses were performed in the individually defined ROIs. **a**. Activity (percentage of signal change, PSC) for facing and non-facing dyads in each ROI. Higher activity for facing (*vs*. non-facing) dyads is taken as marker of configural processing. **b**. Difference in neural activity (PSC) between upright and inverted stimuli (i.e., inversion effect) for facing and non-facing dyads, in each ROI. Greater inversion effect for facing than non-facing dyads is taken as another marker of configural processing. Error bars denote the within-subjects normalized SEM. ^#^*p* = 0.073; \**p* ≤ 0.05; \*\**p* ≤ 0.01; \*\*\**p* ≤ 0.001; All tests are one-tail *t*-tests.

In sum, we found higher activity for facing than non-facing dyads selectively in the EBA, FBA and FFA, that is, in category-specific areas relevant for the processing of the current stimuli. Moreover, neural responses showed a pattern consistent with the behavioral 2BIE (i.e., a larger difference between upright and inverted stimuli for facing than for non-facing dyads) in the FFA and EBA. The relationship between the behavioral and neural BIE was further investigated as follows.

#### Brain-behavior correlation

We investigated whether the behavioral 2BIE, taken as a signature of configural processing of body dyads, could predict the response to dyadic stimuli in visual cortex. This analysis was carried out within the ROIs, using the PSC extracted for each condition, and across the whole-brain. First, for each subject who was included in the behavioral and fMRI sessions of the study, for each ROI, we computed a measure of the neural 2BIE as [(Upright-Inverted)_facing_ / (Upright+Inverted)_facing_] – [(Upright-Inverted)_non-facing_ / (Upright+Inverted)_non-facing_]. Normalized measure of activity was used to take care of inter-subjects differences in the overall magnitude of fMRI response, which are not suitable for correlation with behavioral responses (30). Pearson correlations computed between the individual neural and behavioral 2BIE (accuracy data), revealed a significant effect in the bilateral EBA [*r*(20) = 0.44, p = 0.039], but not in other ROIs [FBA: *r*(20) = 0.16, p > 0.250; FFA: *r*(20) = 0.28, p = 0.214; PPA: *r*(20) = -0.24, p > 0.250; EVC: *r*(20) = 0.27, p = 0.228] (Fig. 5a). For this analysis, we considered the subjects’ performance in the first session of the behavioral task, as that was devoid of learning effects. The same analysis using data from the second session of the behavioral task, showed consistent results, with the strongest effect in the right EBA (see *Supplemental Information 6, 7, 8* for details on correlation analysis using data from the second session, average performance across the two sessions, and for separate left and right ROIs).

**Figure 5.**
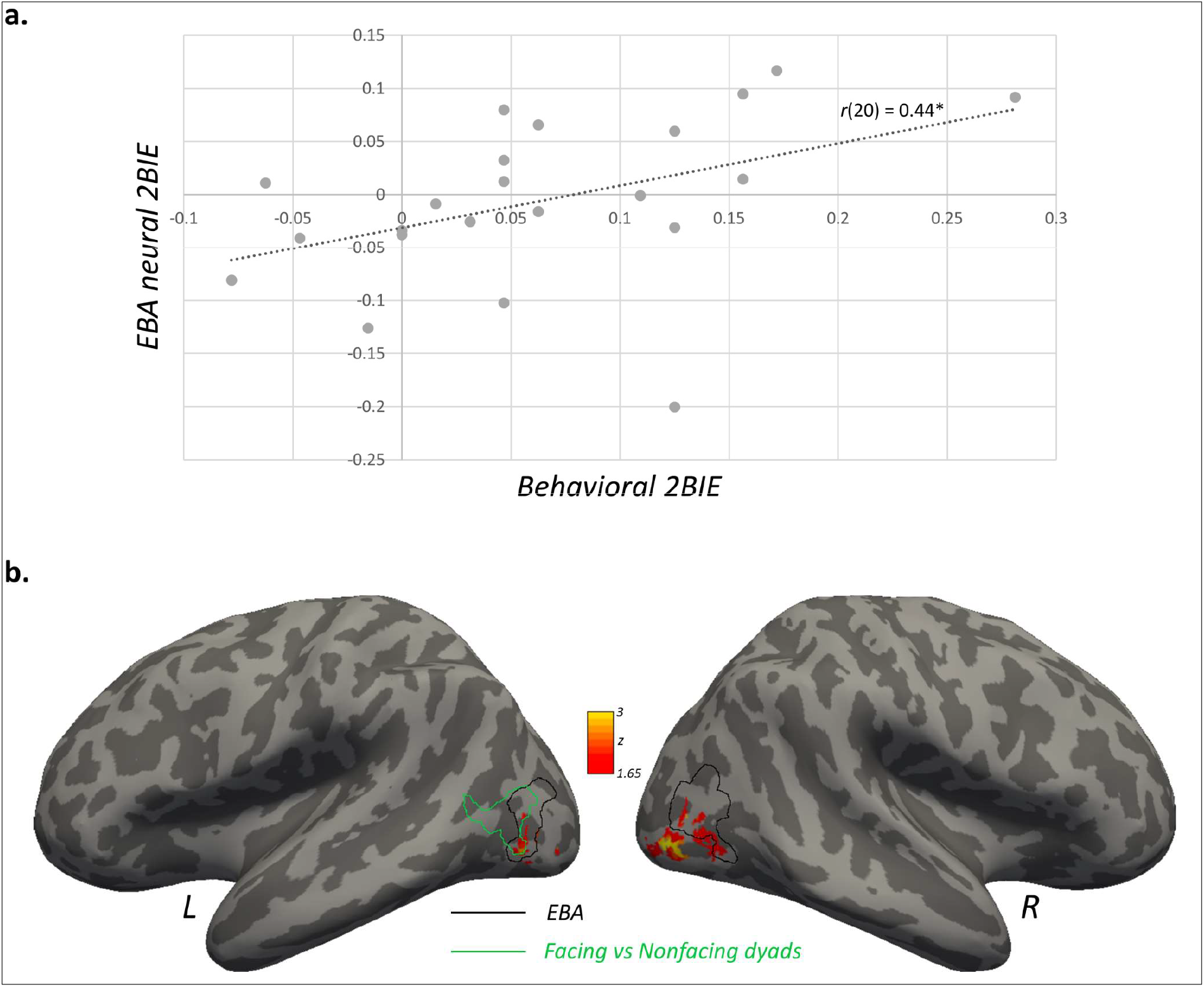
Brain-behavior mapping for the 2BIE. **a**. Correlation between the behavioral 2BIE (average performance in session 1 and 2 of the visual-categorization task) and the neural-2BIE in the functionally defined EBA. \**p* ≤ 0.05 (one-tail). **B**. Statistical map (uncorrected) representing the results of the correlation between the neural 2BIE and the behavioral 2BIE computed in each body-responsive voxel across the whole brain. The color bar indicates *z* values. The region delimited by the black line corresponds to the EBA found with the group-level contrast bodies > [objects+faces+places] using data recorded during the functional localizer task. The region delimited by the green line corresponds to the group-level cluster found with the contrast facing > non-facing dyads in the analysis of data recorded during the main fMRI experiment.

Second, a separate data-driven analysis provided converging evidence for a 2BIE in the EBA. For each subject, for each voxel responsive to body stimuli across the whole brain, we computed the neural 2BIE and performed the correlation with the individual behavioral 2BIE. A significant brain-behavior correlation was found in a bilateral cluster in the lateral occipital cortex (Fig. 5b), encompassing the EBA and extending more posteriorly into the occipital cortex [(right peak MNI coordinates: 44 -82 -6; peak *z*-value = 3.43; peak *p*-value < 0.001; cluster size = 219), (left peak MNI coordinates: -48 -74 2; peak *z*-value = 2.83; peak *p*-value = 0.002; cluster size = 52)]. The effect appeared bilaterally but more strongly in the right hemisphere, where it survived the correction for multiple comparisons (*Supplemental Information 5*).

#### Correlation between the 2BIE and AQ

The above results show that the behavioral 2BIE not only predicts the processing of body dyads in the visual cortex, but also provides a reliable measure of the individual processing of social (multiple-person) scenes, which may be useful to capture interindividual differences. Sensitivity to inversion has been shown to be predictive of the individual’s social abilities (10). To study whether a similar relationship can be established for the 2BIE, all participants in the two behavioral sessions of the fMRI study also completed the Autism Quotient test (AQ), a self-report measure of autistic traits, widely used to quantify the individual’s socially relevant cognitive abilities (31). Correlation between the individual AQ scores and behavioral 2BIE was significant and negative (Spearman *rho*(20) = -0.45; *p* = 0.035), indicating lower sensitivity to dyad inversion in individuals with higher scores for autistic traits (*Supplemental Information 9* for details in correlation analysis using data from the second session and average performance across the two sessions).

### Processing of single bodies

#### The body inversion effect

Configural processing are also thought to apply to single bodies, although the underlying mechanisms seem to be functionally and anatomically distinct from face-related processing (37–39), and, in many aspects, less clear (29, 40). To contribute to this issue, we performed two tests of the body inversion effect as a signature of configural processing. Using fMRI data from runs involving single bodies and non-body objects, we extracted the neural response to upright and inverted stimuli from each ROI. First, in each ROI, we tested whether the difference in the response to upright bodies *vs*. inverted ones was statistically larger than a general effect of inversion that would be observed for any (non-body) stimulus. We ran separate 2 Category x 2 Orientation ANOVAs for each ROI. In the EBA, we found a stronger response to bodies than objects [*F*(1,28) = 160.23, *p* < 0.001, *η*_p_^2^ = 0.85], and to upright than inverted stimuli [*F*(1,28) = 7.81, *p* = 0.009, *η*_p_^2^ = 0.22], but no interaction between Category and Orientation [*F*(1,28) = 0.47, *p* > 0.250, *η*_p_^2^ = 0.02]. Although the interaction was not significant, pairwise *t*-tests (one-tail) showed greater activity for upright *vs*. inverted bodies [*t*(28) = 2.15, *p* = 0.02], and a trend for a greater response to upright *vs*. inverted objects [*t*(28) = 1.56, *p* = 0.065;]. Identical results were found in the FBA [effect of Category: *F*(1,28) = 38.30, *p* < 0.001, *η*_p_^2^ = 0.58; effect of Orientation: *F*(1,28) = 7.53, *p* = 0.010, *η*_p_2 = 0.21; Interaction: *F*(1,28) = 0.24, *p* > 0.250, *η*_p_^2^ = 0.01; upright *vs*. inverted bodies: *t*(28) = 2.30, *p* = 0.015, one-tail; upright *vs*. inverted objects: *t*(28) = 1.45, *p* = 0.079, one-tail] and in the FFA [effect of Category: *F*(1,28) = 41.26, *p* < 0.001, *η*_p_^2^ = 0.60; effect of Orientation: *F*(1,28) = 23.59, *p* < 0.001, *η*_p_^2^ = 0.46; Interaction: *F*(1,28) = 1.30, *p* > 0.250, *η*_p_^2^ = 0.04; upright *vs*. inverted bodies: *t*(28) = 4.21, p < 0.001, one-tail; upright *vs*. inverted objects: *t*(28) = 2.30, p = 0.015, one-tail]. In the PPA, activity was stronger for objects than for bodies [*F*(1,28) = 46.48, *p* < 0.001, *η*_p_^2^ = 0.62], and there was no effect of orientation, [*F*(1,28) = 0.99, *p* > 0.250, *η*_p_^2^ = 0.03], or interaction [*F*(1,28) = 1.93, *p* = 0.176, *η*_p_^2^ = 0.06], no difference between upright and inverted bodies [*t*(28) = 0.29, p > 0.250, one-tail], but a stronger response to upright than inverted objects [*t*(28) = 1.71, *p* = 0.049, one-tail]. In the EVC, we found a trend for a stronger response to objects than bodies [*F*(1,28) = 4.05, *p* = 0.054, *η*_p_^2^ = 0.13], and a stronger response to upright than inverted stimuli [*F*(1,28) = 7.56, *p* = 0.010, *η*_p_^2^ = 0.2], but no interaction [*F*(1,28) = 0.19, *p* > 0.250, *η*_p_^2^ = 0.1], and a stronger response to upright than inverted bodies [*t*(28) = 1.99, *p* = 0.028, one-tail] and to upright than inverted objects [*t*(28) = 2.09, *p* = 0.023, one-tail]. In sum, activity was stronger for upright than inverted bodies in all body- and face-specific ROIs considered here (Fig. 6a), but in none of the ROIs the BIE was statistically different from the effect of inversion on the neural response to non-body objects.

**Figure 6.**
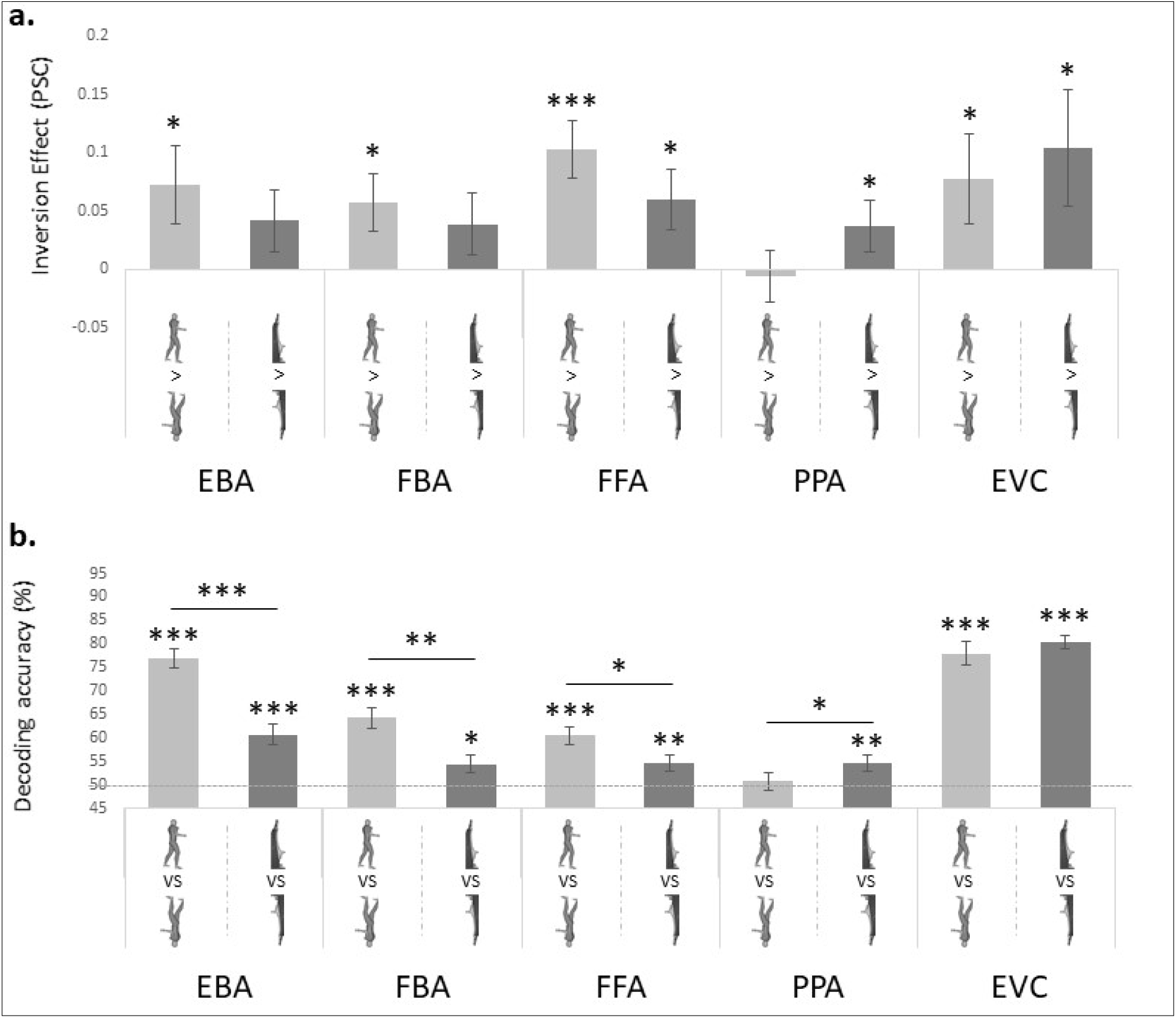
Neural markers of configural body processing in ROIs. **a**. Difference in neural activity (PSC) for upright *vs*. inverted stimuli (i.e., inversion effect) for bodies and objects, in each ROI. In no ROI, the inversion effect was larger for bodies than objects. \**p* ≤ 0.05; \*\*\**p* ≤ 0.001 All tests are one-tail *t*-tests. **b**. Classification accuracies for decoding of upright *vs*. inverted bodies and upright *vs*. inverted objects, in each ROI. Higher deciding accuracy for bodies than objects is taken as a marker of configural processing. Error bars denote the within-subjects normalized SEM. \**p* ≤ 0.05; \*\**p* ≤ 0.01; \*\*\**p* ≤ 0.001; Comparisons against chance level (50 %) are one-tail *t*-tests. Comparisons of accuracies between bodies and objects are two-tail *t*-tests.

Using as another measure of configural body processing, the discrimination accuracy for upright *vs*. inverted bodies in multivariate pattern analysis (MVPA) (29), we found more compelling evidence for the BIE in face- and body-specific ROIs. This analysis showed better decoding of upright, relative to inverted stimuli for both body and non-body objects in the EBA, FBA, FFA, EVC, and for non-body objects in the PPA (Fig. 6b; Table 1). It also showed that discrimination of multivariate patterns of upright *vs*. inverted stimuli was more accurate for bodies than for non-body objects in the EBA, FBA and FFA, was more accurate for non-body objects than bodies in the PPA, and did not differ between the two stimulus classes in the EVC [EBA: *t*(28) = 6.17, *p* < 0.001; FBA: *t*(28) = 3.20, *p* = 0.002; FFA: *t*(28) = 2.05, *p* = 0.025; PPA: *t*(28) = 1.90, *p* = 0.034; EVC: *t*(28) = 0.85, *p* = 0.202]. However, correlations between the individual behavioral BIE and the neural BIE yielded no significant effect in any ROI [EBA: *r*(20) = -0.10, *p* > 0.250; FBA: *r*(20) = -0.05, *p* > 0.250; FFA: *r*(20) = -0.12, *p* > 0.250; PPA: *r*(20) = 0.88, *p* > 0.250; EVC: *r*(20) = 0.07, p > 0.250], or in any voxel responsive to bodies, across the brain.

**Table 1.** Mean classification accuracy (± SEM) and significance for the decoding of upright *vs*. inverted single bodies and upright *vs*. inverted single objects as a multivariate test of the inversion effect. \*\**p* ≤ 0.01; \*\*\**p* ≤ 0.001; All tests are one-tail *t*-tests

| ROI | Bodies |  |  | Objects |  |  |
| --- | --- | --- | --- | --- | --- | --- |
| | Accuracy | $t$ (28) | $p$-value | Accuracy | $t$ (28) | $p$-value |
| EBA | 76.80 (± 2.04) | 13.16 | *** $< 0.001$ | 60.70 (± 2.09) | 5.12 | *** $< 0.001$ |
| FBA | 64.22 (± 2.15) | 6.61 | *** $< 0.001$ | 54.38 (± 1.97) | 2.23 | ** 0.017 |
| FFA | 60.42 (± 2.02) | 5.15 | *** $< 0.001$ | 54.53 (± 1.72) | 2.63 | ** 0.007 |
| PPA | 50.72 (± 1.95) | 0.37 | $> 0.250$ | 54.60 (± 1.61) | 2.85 | ** 0.004 |
| EVC | 78.02 (± 2.62) | 10.70 | *** $< 0.001$ | 80.39 (± 1.56) | 19.44 | *** $< 0.001$ |

## Discussion (1377)

Our findings show that, during processing of multiple bodies, face- and body-specific visual areas encode spatial relations between bodies, yielding different neural responses to face-to-face and back-to-back bodies. In particular, neural responses to facing bodies showed the signatures of configural processing, such as stronger activity relative to the response to their scattered version (non-facing dyads), greater susceptibility to stimulus inversion, and correlation with the behavioral 2BIE, taken as a marker of configural facing-dyad processing (24). Finally, we found that the 2BIE was reliable at the individual level, across behavioral sessions and across brain and behavioral measurement, and correlated with the individual’s social sensitivity coarsely captured by the AQ.

### Configural processing of body dyads

Previous studies have reported increased activity for facing than non-facing dyads in face- and body-specific visual areas ((33, 41, 42). Thus, depending on their relative spatial positioning, the same two bodies are visually processed in different ways. This result is compatible with growing evidence showing that the processing of spatial relations is part of, and affects, the object recognition process (25, 33, 43, 44). In addition, here we showed particularly high sensitivity of the EBA and FFA to the inversion of facing (*vs*. non-facing) dyads, which characterizes the increased neural response to facing dyads, in terms of configural processing.

Although there was only a trend for the interaction between Stimulus (facing/non-facing dyads) and Orientation (upright/inverted) in the EBA, this was the only ROI, in which the activity was predicted by the performance-based 2BIE measured during the visual-categorization task. This result was confirmed by the whole brain-behavior correlation, which tested every voxel responsive to body-stimuli across the brain, and revealed an effect in the lateral occipital cortex. We propose that this effect, encompassing the EBA and peaking in an occipital site between the EBA and a posterior occipital area adjacent or overlapping with another face-specific region (occipital face area; see 45), reveals a new uncharted area of functional specialization for processing spatial relations between bodies, or people.

Like the EBA (in effect, more strongly than the EBA), the FFA showed high sensitivity to the inversion of facing (relative to non-facing) dyads (i.e., Stimulus by Orientation interaction), but no correlation with the behavioral 2BIE. One possibility is that the FFA response is tied to the processing of faces (and their spatial relations), which were clearly visible during fMRI, and less visible, or less attended to, in the behavioral task –in the visual-categorization task, subjects were instructed to recognize bodies, which varied in posture but always had the same sketchy head/face (see Fig. 2). A version of the visual-categorization task with more emphasis on faces and face processing might yield brain-behavioral correlation in the FFA. This reasoning implies that the visual processing of spatial relations would be to some extent category-specific, involving the same areas that respond preferentially to the object class implicated in the relation: body-specific areas for bodies, and face-specific areas for faces. However, the recruitment of the FFA in a task that emphasizes body processing is also compatible with the view that the FFA is involved in configural analysis of both faces and bodies (29; see also 46–48). On this view, different response profiles in the FFA and EBA may reflect the implementation of different aspects, or kinds, of configural processing (7, 49).

### 2BIE and interindividual differences

Face recognition abilities vary across individuals (50–54), but are fairly consistent at the individual level (55–58), and best predicted by the magnitude of face inversion effect, among the markers of configural face processing (49). Evidence for reduced sensitivity to face inversion in ASD subjects has encouraged the thinking that the face inversion effect can be used as a marker of alterations and disease risks that impact social cognition (8, 11–15).

Along those lines, we showed that, in body-dyad perception, the 2BIE varies across individuals, is reliable over time at the individual level, and correlates with a physiological measure of the same processing (i.e., activity in visual areas for face/body processing; see 59). Moreover, we found that the 2BIE correlates with autism-spectrum traits, as measured with the AQ test, so that the magnitude of the 2BIE decreases as the AQ score increases. The relationship between visual configural processing for body-dyads –as well as for faces and biological motion (10, 60)– and the set of abilities measured with the AQ, remains unclear. While we work toward explaining this relationship, effects that reliably characterize the individual’s performance, highlighting population-level and interindividual differences, can offer tools for detection of diseases or vulnerabilities. In this spirit, our results encourage further investigation on the clinical significance of the (behavioral and neural) 2BIE as a biomarker of socially relevant visuoperceptual functions.

### Body dyads and single bodies

Targeted as a measure of configural processing, the body inversion effect (BIE) has been demonstrated with behavioral tasks analogous to those used in face perception (22, 61), and effects in the electroencephalography signal (62–64). However, different fMRI measures of the BIE (e.g., neural adaptation or MVPA) have yielded only inconsistent effects of body inversion in the FFA and/or in the EBA (29, 40), and some studies have ascribed those effects to the processing of head/face information rather than whole body (40, 65). Thus, it remains debated whether bodies evoke reliable inversion effect, or recruit configural processing at all.

Our results confirm the ambiguity of the BIE. The behavioral measure of the BIE showed a cost of inversion that was smaller than the cost for facing dyads (and comparable to the cost for non-facing dyads; Fig. 2). Moreover, the effect of body inversion in body- and face-specific visual areas was statistically different from a general effect of inversion (i.e., inversion of non-body objects) in the multivariate test (discrimination of upright *vs*. inverted stimuli), but not in the univariate test (level of activity for upright minus inverted bodies) and, in none of the ROIs, it correlated with the behavioral BIE.

While leaving open the question of the origin of the BIE and its relationship with the 2BIE, our findings suggest that the two effects differ in kind and/or degree of underlying mechanisms. Moreover, we note that the behavioral 2BIE was correlated with neural activity that partly –but not entirely– overlapped with the EBA, as defined with the classic contrast [single bodies > non-body objects]. This observation encourages the hypothesis that the visual processing of body dyads recruits additional or partly different mechanisms (and neural structures) with respect to the visual processing of single bodies.

## Conclusions

Using a multimodal approach, we connected behavioral (the 2BIE) and neural phenomena (greater sensitivity to inversion for facing *vs*. non-facing dyads and larger response to facing *vs*. non-facing dyads), which together demonstrate the recruitment of specialized configural processing in visual perception of minimal social scenes (i.e., facing-body dyads). The recruitment of configural processing suggests that facing dyads are processed with the same efficiency and high specialization of other highly familiar and/or biologically relevant classes of stimuli, such as faces, bodies and biological motion. Our investigation on the 2BIE also shows that this effect matches –i.e., can account for– the neural effects of body dyad perception, is reliable at the individual level, and correlates with an individual’s AQ. The current characterization of the 2BIE thus supports its value as a marker of configural processing and, possibly, a biomarker of individual visuoperceptual abilities. The visuoperceptual abilities captured by the 2BIE could be precisely those that lay the foundations for living in a social world, for our ability to understand others’ relationships and interactions.

## Material and Methods

### Participants

Thirty subjects took part in the fMRI study (16 female; mean age 24.8 years, *SD* = 4.6). All had normal or corrected-to-normal vision and reported no history of psychiatric or neurological disorders, or use of psychoactive medications. They were screened for contraindications to fMRI and gave informed consent before participation. Of the 30 subjects who enrolled in the fMRI experiment, 23 agreed to come back twice, to take part in two sessions of the behavioral study (13 females; mean age 24.6 years ± 4.5 *SD*). This number was comparable to the sample size used in previous research that used similar task and design (24). From behavioral and fMRI data analyses, we excluded one subject for whom average performance in the behavioral task was >3 *SD* above the group mean. Thus, the final fMRI analyses included 29 participants, and the final behavioral analyses included 22 participants. This study was approved by the local ethics committee (CPP Sud Est V, CHU de Grenoble).

### Stimuli

#### Behavioral study

Sixteen grayscale renderings of single human bodies in profile view and various biomechanically possible poses, were created and edited with Daz3D (Daz Productions, Salt Lake City) and the Image Processing Toolbox of MATLAB (The MathWorks Inc, Natick, Massachusetts). As many bodies were obtained by flipping horizontally each unique body, which yielded to a total of 32 single bodies. Sixteen facing dyads were created from the 32 single bodies, and then they were horizontally flipped to create 16 new dyads, for a total of 32 facing dyads. Non-facing dyads were created by swapping the position of the two bodies in each facing dyad (i.e., the body on the left side was moved to the right side and *vice versa*). The distance between bodies (i.e., the distance between the two closest points of the two bodies in a dyad) was matched across facing and non-facing stimuli (mean_facing_ = 82.88 pixels ± 13.76 *SD*; mean_non-facing_ = 83.06 ± 13.86 *SD*; *t*(15) = 1.00; *p* > 0.250). For single-body stimuli, the center of image corresponded to the center of the body, defined as the mid-point between the most extreme point on the left and on the right along the x-axis. For dyads, the center of the image was between, at equal distance from, the centers of the two bounding boxes that contained each body. In summary, the final stimulus set included 96 stimuli: 32 single bodies, 32 facing dyads and 32 non-facing dyads, differing only for the relative spatial positioning of the two bodies. Another set of stimuli included 96 images of single chairs (32) and pairs of chairs (32 facing and 32 non-facing). Chair-stimuli were created from 16 grayscale exemplars of chairs and their horizontally flipped version, which were combined in 16 pairs of facing pairs and their flipped version, and 16 pairs of non-facing chairs and their flipped version. All body- and chair-stimuli were inverted upside-down, yielding a total of 384 stimuli presented against a light grey background. The same number of masking stimuli was created, consisting of high-contrast Mondrian arrays (11° x 10°) of grayscale circles of variable diameter (0.4°-1.8°).

#### fMRI study

The fMRI study involved the 192 (upright and inverted) body-stimuli used in the behavioral study (64 single bodies, 64 facing dyads and 64 non-facing dyads), in addition to the same 32 single chairs and a new set of 32 machines. Images in the last set depicted one of eight electronic devices (four automated teller machine, and four game machines; i.e., slot machines or arcade game machines) presented in lateral view upright (N=8) and inverted (N=8), and their horizontally flipped version, upright (N=8) and inverted (N=8).

### Procedures

#### Behavioral study

##### Visual categorization task

The task included two identical runs, each containing 32 trials for each of the twelve conditions (upright and inverted single, facing and non-facing, bodies and chairs), presented in random order. Each stimulus appeared once in a run. Subjects sat on a chair, 60 cm away from a computer screen, with their eyes aligned to the center of the screen (17-in. CRT monitor; 1024×768 pixel resolution; 60-Hz refresh rate). Stimuli on the screen did not exceed 7° of visual angle. Each trial included the following sequence of events: blank screen (200 ms), fixation cross (500 ms), blank screen (200 ms), target stimulus (30 ms), mask (250 ms) and a final blank screen that remained until the subject gave a response. The next trial began after a variable interval between 500 and 1000 ms. For each trial, subjects had to report whether they had seen bodies or chairs, regardless of the number of items (one/two), positioning (facing/non-facing) or orientation (upright/inverted). They responded by pressing one of two keys on a keyboard: “1” with the index finger for “bodies”, or “2” with the middle finger for “chair” or *vice versa* (stimulus-response mapping was counterbalanced across participants). Every 32 trials subjects could take a break. Two blocks of familiarization preceded the experiment. In the first block, stimuli (four per condition) were shown for 250 ms, so that the subjects could easily see them. The second block (eight trials per condition) was identical to the actual experiment. Instructions for the familiarization blocks were identical to those of the actual experiment. The experiment lasted ∼40 min. Stimulus presentation and response collection (accuracy and RTs) were controlled with Psychtoolbox (66) through MATLAB. Each participant took part in two sessions of the same task. The second session took place with a delay ranging from two weeks to one month after the first one (mean interval 19.8 days ± 6.1 *SD*).

##### Autistic-Spectrum Quotient measure

After the second behavioral session, subjects completed the 50-items Autism-Spectrum Quotient scale (AQ; see 31), which measures the degree to which an adult with normal intelligence shows traits associated with the autism spectrum, in the domain of social skills, attention-switching, attention to detail, communication, and imagination. Subjects responded to each item using a 4-point rating scale, ranging from “definitely agree” to “definitely disagree”.

#### fMRI study

##### fMRI data acquisition

Imaging was conducted on a MAGNETOM Prisma 3T scanner (Siemens Healthcare). T2*-weighted functional volumes were acquired using a gradient-echo echo-planar imaging sequence (GRE-EPI; TR/TE = 2000/30 ms, flip angle = 80°, acquisition matrix = 96 × 92, FOV = 210 × 201, 56 transverse slices, slice thickness = 2.2mm, no gap, multiband acceleration factor = 2 and phase encoding set to anterior/posterior direction). For the main experiment and the functional localizer task, we acquired eight runs for a total of 1546 frames per participant. Acquisition of high-resolution T1-weighted anatomical images was performed after the third functional run of the main experiment and lasted 8 min (MPRAGE; TR/TE/TI = 3000/3.7/1100 ms, flip angle = 8°, acquisition matrix = 320 × 280, FOV = 256 × 224 mm, slice thickness = 0.8mm, 224 sagittal slices, GRAPPA accelerator factor = 2). Acquisition of two field maps was performed at the beginning of the fMRI session.

##### Main fMRI experiment

The fMRI data acquisition consisted of two parts: the main fMRI experiment, and a functional localizer task, which we describe below. In the main experiment, facing and non-facing dyads in upright or inverted orientation were presented over three runs, each lasting 6.83 min. Each run consisted of two sequences of 16 blocks (four blocks by condition), for a total of 32 blocks of eight second each. Blocks in the first sequence were presented in a random order, and blocks in the second sequence were presented in the counterbalanced (i.e., reversed) order relative to the first sequence. Each block featured eight images of the same condition, presented in a random order. A stimulus appeared once in a block, and twice in a run (once in each sequence). Thus, each of the three runs included eight blocks per condition (24 across the whole experiment). Three additional runs of 6.83 min each featured images of single bodies and single objects presented upright or inverted. Each run included 16 blocks (eight seconds each), eight with bodies and eight with objects (four with chairs and four with machines). Each block included eight different stimuli of the same condition. Each stimulus appeared twice in a run (once in each sequence). Runs with dyads and runs with single items (bodies or objects) were presented in pseudorandom order to avoid the occurrence of more than two consecutive runs of the same stimulus group. Each run began with a warm-up block (10 s) and ended with a cool-down block (16 s), during which a central fixation cross was presented. Within a run, the onset time of each block was jittered (range of inter-block interval duration: 2-6 s; total inter-block time by runs: 128 s), using the optseq tool of Freesurfer (67) for optimal jittering. During each block, a black cross was always present in the center of the screen, while stimuli appeared for 550 ms, separated by an interval of 450 ms. In the 37% of all the stimulation and fixation periods, the cross changed color (from black to red). Participants were instructed to fixate the cross throughout the experiment, detect and report the color change by pressing a button with their right index finger. This task was used to minimize eye movements and maintain vigilance in the scanner. The main experiment lasted 41 min. During fMRI, stimuli were back-projected onto a screen by a liquid crystal projector (frame rate: 60 Hz; screen resolution: 1024×768 pixels, screen size: 40×30 cm). For all the stimuli, the center of the image overlapped with the center of the screen. Subjects, lying down inside the scanner, viewed the stimuli binocularly (∼7° of visual angle) through a mirror above their head. Stimulus presentation, response collection and synchronization with the scanner were controlled with the Psychtoolbox (66) through MATLAB (The MathWorks Inc, Natick, Massachusetts).

##### Functional localizer task

In addition to the six experimental runs, subjects completed a functional localizer task, with stimuli and task adapted from the fLoc package (68). During this task, subjects saw 180 grayscale photographs of the following five object classes: 1) body-stimuli (headless bodies in various views and poses, and body parts); 2) faces (adults and children); 3) places (houses and corridors); 4) inanimate objects (various exemplars of cars and musical instruments); 5) scrambled objects. Stimuli were presented over two runs (5.27 min each). Each run began with a warm-up block (12 s) and ended with a cool-down block (16 s), and included 72 blocks of four seconds each: 12 blocks for each object class with eight images per block (500 ms per image without interruption), randomly interleaved with 12 baseline blocks featuring an empty screen. To minimize low-level differences across categories, the view, size, and retinal position of the images varied across trials, and each item was overlaid on a 10.5° phase-scrambled background generated from another image of the set, randomly selected. During blocks containing images, some images were repeated twice, interleaved by a different image (2-back task). Participants had to press a button when they detected the repetition.

##### Preprocessing of fMRI data

Functional images were preprocessed and analyzed using MATLAB, in combination with SPM12 (69), the CoSMoMVPA toolbox (70), the MarsBaR toolbox (71) and the MatlabTFCE toolbox (72). The first four volumes of each run were discarded, taking into account initial scanner gradient stabilization (73). Preprocessing of the remaining volumes involved despiking, slice time correction, geometric distortions correction using field maps, spatial realignment and motion correction using the first volume of each run as reference. The maximum displacement after despiking was 0.74 mm on the *x*-axis (mean_max_ = 0.30, *SD* = 0.18), 2.31 mm on the *y*-axis (mean_max_ = 0.65, *SD* = 0.52) and 3.29 mm on the *z*-axis (mean_max_ = 1.06, *SD* = 0.64), which did not exceed the Gaussian kernel of 5 mm FWHM used for the spatial smoothing before estimating the realignment parameters (74). Anatomical volumes were co-registered to the mean functional image, segmented into gray matter, white matter and cerebrospinal fluid in native space, and aligned to the probability maps in the Montreal Neurological Institute (MNI). The DARTEL method (75) was used to create a flow field for each subject and an inter-subject template, which was registered in the MNI space and used for normalization of functional images. Final steps included spatial smoothing with a Gaussian kernel of 6 mm FWHM and removing low-frequency drifts with a temporal high-pass filter (cutoff 128 s).

### Analyses

#### Behavioral study

##### Visual categorization task

Separately for the two sessions of the task, we computed the mean proportion of correct responses (accuracy) and the mean RTs for each condition, for each subject. Mean RTs were computed over trials with accurate response and RTs within 2 *SD* from the individual mean (discarded trials in session 1: 13.10% ±7.21 *SD*; in session 2: 10.20% ±5.87 *SD*). Accuracy and RT data were analyzed with 2 Category (body, chair) x 3 Stimulus (facing dyads, non-facing dyads, single bodies) x 2 Orientation (upright, inverted) repeated-measures ANOVAs. Critical comparisons were tested using pairwise *t* tests (two-tail). Analyses were repeated using average performance values from the two sessions, and for each session separately.

##### Test-retest reliability

For each subject, we computed the 2BIE on accuracy values [(Upright-inverted)_facing_-(Upright-inverted)_non-facing_] for each of the two sessions and compared them (*t* test, α = 0.05, two-tail), to test for difference in the magnitude of the effect between sessions. Second, we computed the within-subjects Pearson correlation (α = 0.05) between the 2BIE values in the first and second session, to assess the intra-individual consistency of the 2BIE. We ran a permutation test (10000 iterations) to assess whether the within-subjects correlation was higher than the correlation between any two randomly selected values (between-subjects correlations). To do so, in each iteration, a subject’s 2BIE from the first session was randomly assigned to the 2BIE of another subject in the second session and a Pearson correlation coefficient was computed. We ranked the single within-subjects correlation value on the distribution of the 10000 between-subjects correlations values, and divided the rank by number of permutations to obtain a *p* value. We assessed whether a significant within-subject correlation was statistically higher than random between-subject correlations.

##### Correlation between 2BIE and AQ

We computed Spearman correlation (α = 0.05) between the behavioral 2BIE and the AQ score of the 22 subjects who participated in both behavioral sessions. This analysis was repeated using data from session 1 of the visual categorization task (*Results*), from session 2 of the task and the average performance values from the two sessions (*Supplementary information*).

#### fMRI study of body dyad processing

##### Whole-brain analysis

For each subject, the blood-oxygen-level-dependent (BOLD) signal of each voxel was estimated with a random-effects general linear model (RFX GLM), including four regressors for the four experimental conditions (upright and inverted facing and non-facing dyads), one regressor for fixation blocks, and six regressors for movement correction parameters as nuisance covariates. In the first analysis, the effect of the visuo-spatial configuration of bodies within a dyad was assessed, separately for the upright and inverted orientations, with a RFX GLM contrasting facing *vs*. non-facing dyads. In the second analysis, the effect of stimuli orientation was assessed, separately for facing and non-facing dyadic configurations, with a RFX GLM contrasting upright > inverted orientation. For all univariate analyses, the statistical significance of second-level (group) effects was determined using a voxelwise threshold of *p* ≤ 0.001, family-wise error corrected at the cluster level.

##### Definition of regions of interest (ROIs)

Individual data recorded during the functional localizer task were entered into a General Linear Model with five regressors for the five object-class conditions (bodies, faces, places, objects and scrambled objects), one regressor for baseline blocks, and six regressors for movement correction parameters as nuisance covariates. Three bilateral masks of the middle temporo-occipital gyrus (MTOG), the temporo-occipital fusiform cortex (TOFC) and the inferior parahippocampal cortex (PHC) were created using FSLeyes (76) and the Harvard-Oxford Atlas (77) through FSL (78). In addition, a bilateral mask of the EVC encompassing visual areas V1/V2 and V3v/V3d, was created using the SPM Anatomy toolbox (79). Within each mask of each subject, we selected the voxels with significant activity (threshold: *p* = 0.05) for the contrasts of interest, to create ROIs: EBA in the MTOG with the contrast bodies > [objects+faces+places], FFA in the TOFC with the contrast faces > [objects+bodies+places], FBA in the TOFC with the contrast body > [objects+face+places] and PPA in the PHC with the contrast places > [objects+faces+bodies]. The EVC-ROI included all the voxels responsive to visual stimuli (contrast: [bodies+faces+places+objects+scrambled objects] > baseline). For each ROI, all of the voxels within the bilateral mask that passed the threshold were ranked by activation level based on *t* values. The final ROIs included up to 200 best voxels across the right and left ROI (number of voxels in the EBA, FBA, FFA and EVC: 200; in the PPA: 199.86 ± 0.58 *SD*). Neural activity values used for all ROIs analyses were extracted as the percent signal change (PSC) for each condition, using the MarsBaR toolbox (Brett et. al., 2002). When a functional ROI could not be identified (the FBA for one subject and all five ROIs for another, due to a technical failure during the functional localizer task), PSC were extracted from a mask corresponding to the group-level activation for the contrast of interest.

##### ROIs analyses

Using data recorded during dyad runs of the main fMRI experiment (N = 29), we analyzed in each ROI the effects of seeing upright or inverted facing and non-facing dyads. In particular, we performed a repeated-measures ANOVA with factors: 5 ROI (EBA, FBA, FFA, PPA and EVC) x 2 Stimulus (facing and non-facing dyads) x 2 Orientation (upright and inverted). Next, pairwise *t*-tests were used to investigate, in each ROI, the two signatures of configural processing: 1) larger response to facing than to non-facing (upright) dyads (α = 0.05 one-tail), and 2) larger difference between upright and inverted stimuli for facing, than for non-facing dyads, or the neural 2BIE (α = 0.05, one-tail).

##### Brain-behavior correlation (in ROIs)

For the 22 subjects who participated in both the behavioral and fMRI sessions, we computed the (normalized) neural 2BIE, using the PSC extracted from each ROI for each condition (upright and inverted facing and non-facing dyads). For each ROI, the values representing the individual neural 2BIEs were correlated with the values representing the individual behavioral 2BIE (Pearson correlation, α = 0.05). This analysis was repeated using data from session 1 of the behavioral task (*Results*), from session 2 and the average performance across the two sessions (*Supplementary Information 7,8*).

##### Brain-behavior correlation (across the whole-brain)

For the 22 participants who participated in both the behavioral and the fMRI sessions, we used the behavioral-2BIE from the first behavioral session and we measured a neural 2BIE for each voxel in the brain. To do so, we selected all the voxels across the brain with a positive β-weight for all conditions of body dyads and, for the selected voxels, we extracted the β-weights for each condition. Then, separately for facing and non-facing dyads, we computed for each voxel the normalized inversion effects ([upright stimuli – inverted stimuli] / (upright stimuli + inverted stimuli]) and we defined the neural-2BIEs as the difference in the normalize inversion effect between facing and non-facing dyads. For all participants, for each voxel, we computed the Pearson correlation between the neural-2BIE and the behavioral-2BIE. For this analysis, the statistical significance of the correlation was determined using a voxelwise threshold of p ≤ 0.05 (one-tailed). The statistical map of Pearson correlation coefficients (Fisher-transformed) was corrected for multiple comparisons using a permutation testing approach (80) combined with threshold free cluster enhancement transformations (TFCE, 81) through the MatlabTFCE toolbox (permutation number = 5000, H = 2, E = 0.5, dh=0.1, Connectivity = 26, one-tailed). The same analysis was repeated using values of the behavioral-2BIE based on session 2 of the behavioral task and on the average of the two sessions (see *Supplementary information 7,8* for results on these analyses).

##### Test of the single-body inversion effect (BIE)

To assess the recruitment of configural processing dfor single bodies, we tested whether the difference in the response to upright bodies *vs*. inverted ones was statistically larger than a general effect of inversion that would be observed for any (non-body) stimulus. For each of the above ROIs, data recorded during the runs of the main fMRI experiment involving single bodies and objects were analyzed with a repeated-measures ANOVA with factors, 2 Category (body and object) and 2 Orientation (upright and inverted). Next, pairwise *t*-tests were used to investigate, in each ROI, the inversion effect for single bodies. In particular, we first computed the inversion effect (stronger response to upright than inverted stimuli) separately for bodies and objects (α = 0.05, one-tail), and then tested whether the inversion effect was larger for bodies than for objects (α = 0.05 one-tail). Moreover, as another test of the body inversion effect, from each ROI, we extracted multivariate activity patterns for the four conditions, and performed MVPA to compute the accuracy of discrimination between upright and inverted stimuli, for bodies and objects. A larger inversion effect for bodies relative to any other object should yield higher discrimination accuracy between upright and inverted bodies, than between upright and inverted (non-body) objects (29). To do so, a support vector machine classifier was trained on 5 out of 6 runs to classify upright *vs*. inverted stimuli, and then tested on data from the remaining run. This scheme was iterated until all runs were used as testing and training sets. For each subject, for each ROI, accuracy values were averaged across all iterations and tested against chance (50% accuracy, *t* tests, α = 0.05, one-tail) and, finally, compared between bodies and objects (*t* tests, α = 0.05, one-tail).

## Data availability statement

Data and codes for the main analyses are available here: https://osf.io/pse3k/

## Acknowledgements

This work was supported by a European Research Council Starting Grant to L.P. (Grant number: THEMPO-758473).

We thank Nicolas Goupil for contributing to the analyses and editing of Figure 1.

