## supplementary_materials for "A new behavioral and neural marker of social vision"

### Supplementary information

**1. Accuracy and RTs results of the behavioral task (session 1).** Accuracy data for the first session of the task were analyzed in a 2 Category (body, chair) x 3 Stimulus (facing, non-facing, single stimuli) x 2 orientation (upright, inverted) repeated-measures ANOVA. Results showed main effects of Category [$F(1,21) = 4.70$, $p = 0.042$, $\eta_p^2 = 0.18$], Stimulus [$F(2,42) = 4.17$, $p = 0.022$, $\eta_p^2 = 0.17$] and Orientation [$F(1,21) = 21.27$, $p < 0.001$, $\eta_p^2 = 0.50$], and significant interactions between Category and Stimulus [$F(2,42) = 6.75$, $p = 0.003$, $\eta_p^2 = 0.24$], Category and Orientation [$F(1,21) = 8.87$, $p = 0.007$, $\eta_p^2 = 0.30$] and between Stimulus and Orientation [$F(2,42) = 6.13$, $p = 0.005$, $\eta_p^2 = 0.23$]. Main effects and interactions were qualified by a three-way interaction between Category, Stimulus and Orientation [$F(2,42) = 4.45$, $p = 0.018$, $\eta_p^2 = 0.17$]. Next, two separate ANOVAs with body-trials only and chair-trials only, showed that the magnitude of the inversion effect was affected by the type of Stimulus (single, facing, non-facing), when stimuli were bodies (Stimulus: [$F(2,42) = 7.52$, $p = 0.002$, $\eta_p^2 = 0.26$]; Orientation: [$F(1,21) = 15.05$, $p < 0.001$, $\eta_p^2 = 0.42$], Interaction: [$F(2,42) = 8.74$, $p < 0.001$, $\eta_p^2 = 0.29$]), but not chairs (Stimulus: [$F(2,42) = 0.95$, $p > 0.250$, $\eta_p^2 = 0.04$]; Orientation: [$F(1,21) = 1.24$, $p > 0.250$, $\eta_p^2 = 0.06$], Interaction: [$F(2,42) = 0.97$, $p > 0.250$, $\eta_p^2 = 0.04$]). Pairwise *t* tests (Fig. S1a) showed that, while the inversion effect was significant for all three body stimulus conditions [facing bodies: $t(21) = 4.33$, $p < 0.001$; non-facing bodies: $t(21) = 3.34$, $p = 0.003$; single bodies: $t(21) = 3.48$, $p = 0.002$], it was larger for facing bodies relative to non-facing bodies [$t(21) = 3.47$, $p = 0.002$] and single bodies [$t(21) = 3.35$, $p = 0.003$], and comparable for the last two [$t(21) = 0.39$, $p > 0.250$]. Pairwise *t* tests for chairs showed an inversion effect for facing and single chairs but not for non-facing chairs [facing chairs: $t(21) = 3.05$, $p = 0.006$; non-facing chairs: $t(21) = 1.22$, $p = 0.234$; single chairs: $t(21) = 2.13$, $p = 0.045$]. There was no significant difference in the magnitude of the inversion effect across the three conditions [facing *vs.* non-facing chairs: $t(21) = 0.98$, $p > 0.250$; single *vs.* facing chairs: $t(21) = 0.93$, $p = 0.192$; single *vs.* non-facing chairs: $t(21) = 0.09$, $p > 0.250$].

RT data for the first session of the task were analyzed in a 2 Category (body, chair) x 3 Stimulus (facing, non-facing, single stimuli) x 2 orientation (upright, inverted) repeated-measures ANOVA. Results showed main effects of Category [$F(1,21) = 26.71$, $p < 0.001$, $\eta_p^2 = 0.56$], a trend for an effect of Stimulus [$F(2,42) = 3.04$, $p = 0.058$, $\eta_p^2 = 0.13$] and an effect of Orientation [$F(1,21) = 49.50$, $p < 0.001$, $\eta_p^2 = 0.70$], and no interaction between Category and Stimulus [$F(2,42) = 0.24$, $p > 0.250$, $\eta_p^2 = 0.01$], an interaction between Category and Orientation [$F(1,21) = 8.35$, $p = 0.009$, $\eta_p^2 = 0.28$] and between Stimulus and Orientation [$F(2,42) = 3.98$, $p = 0.026$, $\eta_p^2 = 0.16$]. The interaction between Category, Stimulus and Orientation was not significant [$F(2,42) = 0.27$, $p > 0.250$, $\eta_p^2 = 0.01$]. Next, two separate ANOVAs with body-trials only and chair-trials only, showed that the magnitude of the inversion effect was affected by the type of Stimulus (single, facing, non-facing), when stimuli were bodies (Stimulus: [$F(2,42) = 1.71$, $p = 0.193$, $\eta_p^2 = 0.08$]; Orientation: [$F(1,21) = 37.82$, $p < 0.001$, $\eta_p^2 = 0.64$], Interaction: [$F(2,42) = 3.46$, $p = 0.041$, $\eta_p^2 = 0.14$]), but not chairs (Stimulus: [$F(2,42) = 1.97$, $p > 0.152$, $\eta_p^2 = 0.09$]; Orientation: [$F(1,21) = 28.34$, $p < 0.001$, $\eta_p^2 = 0.57$], Interaction: [$F(2,42) = 1.41$, $p > 0.250$, $\eta_p^2 = 0.06$]). Pairwise *t* tests (Fig. S1b) showed that, while the inversion effect was significant for all three body stimulus conditions [facing bodies: $t(21) = 4.33$, $p < 0.001$; non-facing bodies: $t(21) = 3.34$, $p = 0.003$; single bodies: $t(21) = 3.48$, $p = 0.002$], it was larger for facing bodies relative to non-facing bodies [$t(21) = 3.47$, $p = 0.002$] and single bodies [$t(21) = 3.35$, $p = 0.003$], and comparable for the last two [$t(21) = 0.39$, $p > 0.250$]. Pairwise *t* tests for chairs showed an inversion effect for facing and single chairs but not for non-facing chairs [facing chairs IE: $t(21) = 3.05$, $p = 0.006$; non-facing chairs IE: $t(21) = 1.22$, $p = 0.234$; single chairs IE: $t(21) = 2.13$, $p = 0.045$]. However, there was no significant difference in the inversion effect across the three conditions [facing chairs *vs.* non-facing chairs: $t(21) = 0.98$, $p > 0.250$; single chairs *vs.* facing chairs: $t(21) = 0.93$, $p = 0.192$; single chairs *vs.* non-facing chairs: $t(21) = 0.09$, $p > 0.250$].

**2. Accuracy and RTs results in the behavioral task (session 2).** Accuracy data of the second session of the task were analyzed in a 2 Category (body, chair) x 3 Stimulus (facing, non-facing, single stimuli) x 2 orientation (upright, inverted) repeated-measures ANOVA. Results showed no main effect of Category [$F(1,21) = 0.96$, $p > 0.250$, $\eta_p^2 = 0.04$] but main effects of Stimulus [$F(2,42) = 4.76$, $p = 0.014$, $\eta_p^2 = 0.18$] and Orientation [$F(1,21) = 16.90$, $p < 0.001$, $\eta_p^2 = 0.45$], and significant interactions between Category and Stimulus [$F(2,42) = 6.24$, $p = 0.004$, $\eta_p^2 = 0.23$], Category and Orientation [$F(1,21) = 14.89$, $p < 0.001$, $\eta_p^2 = 0.41$] and Stimulus and Orientation [$F(2,42) = 3.96$, $p = 0.027$, $\eta_p^2 = 0.16$]. Main effects and interactions were qualified by a three-way interaction between Category, Stimulus and Orientation [$F(2,42) = 7.90$, $p = 0.001$, $\eta_p^2 = 0.27$]. Next, two separate ANOVAs with body-trials only and chair-trials only, showed that the magnitude of the inversion effect was affected by the type of

Stimulus (single, facing or non-facing), for body-stimuli (Stimulus: [$F(2,42) = 6.89$, $p = 0.003$, $\eta_p^2 = 0.24$]; Orientation: [$F(1,21) = 18.00$, $p < 0.001$, $\eta_p^2 = 0.46$], Interaction: [$F(2,42) = 5.78$, $p = 0.006$, $\eta_p^2 = 0.22$]), and chair-stimuli (Stimulus: [$F(2,42) = 0.46$, $p > 0.250$, $\eta_p^2 = 0.02$]; Orientation: [$F(1,21) = 1.83$, $p = 0.190$, $\eta_p^2 = 0.08$], Interaction: [$F(2,42) = 5.96$, $p = 0.005$, $\eta_p^2 = 0.22$]). Pairwise *t* tests (Fig. S1a) showed that, while the inversion effect was significant for all three body conditions [facing bodies: $t(21) = 4.27$, $p < 0.001$; non-facing bodies: $t(21) = 4.15$, $p < 0.001$; single bodies: $t(21) = 3.64$, $p = 0.001$], it was larger for facing bodies relative to non-facing bodies [$t(21) = 2.87$, $p = 0.009$] and single bodies [$t(21) = 2.13$, $p = 0.045$], and comparable for the last two [$t(21) = 1.57$, $p =$ 0.131]. Pairwise *t* tests for chair-stimuli showed an inversion effect for single chairs but not for facing and non-facing chairs [facing chairs IE: $t(21) = 0.50$, $p > 0.250$; nonfacing chairs IE: $t(21) = 0.23$, $p > 0.250$; single chairs IE: $t(21) = 2.80$, $p = 0.010$]. The inversion effect was larger for single chairs relative to facing chairs [$t(21) = 2.64$, $p = 0.015$], and to non-facing chairs [$t(21) = 2.73$, $p > 0.012$] and did not differ between facing and non-facing chairs [$t(21) = 0.77$, $p > 0.250$].

RTs data for the second session of the task were analyzed in a 2 Category (body, chair) x 3 Stimulus (facing, non-facing, single stimuli) x 2 orientation (upright, inverted) repeated-measures ANOVA. Results showed main effects of Category [$F(1,21) = 11.67$, $p = 0.003$, $\eta_p^2 = 0.37$], Stimulus [$F(2,42) = 4.51$, $p = 0.017$, $\eta_p^2 = 0.18$] and Orientation [$F(1,21) = 59.27$, $p < 0.001$, $\eta_p^2 = 0.74$], no interactions between Category and Stimulus [$F(2,42) = 0.21$, $p > 0.250$, $\eta_p^2 = 0.01$] and Stimulus and Orientation [$F(2,42) = 0.31$, $p > 0.250$, $\eta_p^2 = 0.01$], but an interaction between Category and Orientation [$F(1,21) = 15.53$, $p < 0.001$, $\eta_p^2 = 0.43$]. Main effects and interactions were qualified by a three-way interaction between Category, Stimulus and Orientation [$F(2,42) = 3.32$, $p = 0.046$, $\eta_p^2 = 0.14$]. Next, two separate ANOVAs with body-trials only and chair-trials only, showed that the magnitude of the inversion effect was not affected by the type of Stimulus (single, facing, non-facing), when stimuli were bodies (Stimulus: [$F(2,42) = 2.90$, $p = 0.066$, $\eta_p^2 = 0.12$]; Orientation: [$F(1,21) = 56.71$, $p < 0.001$, $\eta_p^2 = 0.73$], Interaction: [$F(2,42) = 1.83$, $p = 0.172$, $\eta_p^2 = 0.08$]), or chairs (Stimulus: [$F(2,42) = 1.99$, $p = 0.149$, $\eta_p^2 = 0.09$]; Orientation: [$F(1,21) = 4.98$, $p = 0.037$, $\eta_p^2 = 0.19$], Interaction: [$F(2,42) = 0.55$, $p > 0.250$, $\eta_p^2 = 0.03$]). Pairwise *t* tests (Fig. S1b) showed that, the inversion effect was significant for all three body-stimulus conditions [facing bodies: $t(21) = 4.83$, $p < 0.001$; non-facing bodies: $t(21) = 4.83$, $p < 0.001$; single bodies: $t(21) = 8.08$, $p < 0.001$], but it was comparable for facing bodies relative to non-facing bodies [$t(21) = 1.60$, $p = 0.123$] and single bodies [$t(21) = 0.49$, $p = 0.629$], and between non-facing bodies and single bodies [$t(21) = 1.77$, $p = 0.092$]. Pairwise *t* tests for chairs showed an inversion effect for non-facing chairs but not for facing and single chairs [facing chairs: $t(21) = 1.16$, $p > 0.250$; non-facing chairs: $t(21) = 2.59$, $p = 0.017$; single chairs: $t(21) = 1.51$, $p = 0.146$]. There was no significant difference between all chair-stimulus conditions [facing chairs *vs.* non-facing chairs: $t(21) = 1.05$, $p > 0.250$; single chairs *vs.* facing chairs: $t(21) = 0.43$, $p > 0.250$; single chairs *vs.* non-facing chairs: $t(21) = 0.58$, $p > 0.250$].

**3. Accuracy results for chairs stimuli in the behavioral task (average of session 1 and 2).** Pairwise *t* tests using accuracy data of chair-stimuli averaged across the two sessions of the behavioral task showed an inversion effect for single chairs, but not for facing and non-facing chairs [single chairs: $t(21) = 4.27$, $p = 0.004$; facing chairs: $t(21) = 1.67$, $p = 0.109$; non-facing chairs: $t(21) = 0.99$, $p > 0.250$]. Using accuracy data, there was no significant difference between all three stimulus conditions [single chairs *vs.* facing chairs: $t(21) = 1.46$, $p = 0.159$; single chairs *vs.* non-facing chairs: $t(21) = 1.44$, $p = 0.165$; facing chairs *vs.* non-facing chairs: $t(21) = 0.36$, $p > 0.250$].

**4. RTs results of the behavioral task (average of session 1 and 2).**

RTs data averaged across the two sessions of the task were analyzed in a 2 Category (body or chair) x 3 Stimulus (facing, nonfacing or single stimuli) x 2 orientation (upright or inverted) repeated-measures ANOVA. Results showed a main effect of Category [$F(1,21) = 20.65$, $p < 0.001$, $\eta_p^2 = 0.50$], Stimulus [$F(2,42) = 6.93$, $p = 0.003$, $\eta_p^2 = 0.25$] and Orientation [$F(1,21) = 60.29$, $p < 0.001$, $\eta_p^2 = 0.74$], and an interaction between Category and Orientation [$F(1,21) = 15.64$, $p < 0.001$, $\eta_p^2 = 0.43$] but no interaction between Category and Stimulus [$F(2,42) = 0.27$, $p > 0.250$, $\eta_p^2 = 0.01$] and Stimulus and Orientation [$F(2,42) = 1.53$, $p = 0.229$, $\eta_p^2 = 0.07$]. Main effects and interactions were qualified by a trend for a three-way interaction between Category, Stimulus and Orientation which was compatible with the accuracy results but did not approach significance [$F(2,42) = 2.73$, $p = 0.077$, $\eta_p^2 = 0.12$]. Next, two separate ANOVAs with body-trials only and chair-trials only, showed results similar to the accuracy results. The magnitude of the inversion effect was affected by the type of Stimulus (single, facing or non-facing), when stimuli were bodies (Stimulus: [$F(2,42) = 3.46$, $p = 0.041$, $\eta_p^2 = 0.14$]; Orientation: [$F(1,21) = 55.24$, $p < 0.001$, $\eta_p^2 = 0.72$], Interaction:

[$F(2,42) = 2.77$, $p = 0.074$, $\eta_p^2 = 0.12$]), but not chairs (Stimulus: [$F(2,42) = 2.83$, $p = 0.071$, $\eta_p^2 = 0.12$]; Orientation: [$F(1,21) = 18.26$, $p < 0.001$, $\eta_p^2 = 0.47$], Interaction: [$F(2,42) = 0.69$, $p > 0.250$, $\eta_p^2 = 0.03$]). Pairwise *t* tests using RTs data averaged across the two sessions of the behavioral task (Fig. S1c) showed that, while the inversion effect was significant for all three body-stimulus conditions [facing bodies: $t(21) = 5.68$, $p < 0.001$; non-facing bodies: $t(21) = 6.26$, $p < 0.001$; single bodies: $t(21) = 8.59$, $p = 0.001$], the effect was qualitatively larger for facing bodies relative to non-facing bodies [$t(21) = 1.85$, $p = 0.078$] and single bodies [$t(21) = 1.74$, $p = 0.097$], and comparable for the last two stimulus conditions [$t(21) = 0.76$, $p > 0.250$]. Pairwise *t* tests for chair-stimuli using RTs data averaged across the two sessions of the behavioral task showed an inversion effect for all three stimulus conditions [single chairs: $t(21) = 3.64$, $p = 0.002$; facing chairs: $t(21) = 4.27$, $p < 0.001$; non-facing chairs: $t(21) = 2.58$, $p = 0.018$]. with no difference across conditions [single chairs *vs.* facing chairs: $t(21) = 1.35$, $p = 0.192$; single chairs *vs.* non-facing chairs: $t(21) = 0.85$, $p = 0.405$; facing chairs *vs.* non-facing chairs: $t(21) = 0.10$, $p > 0.250$]

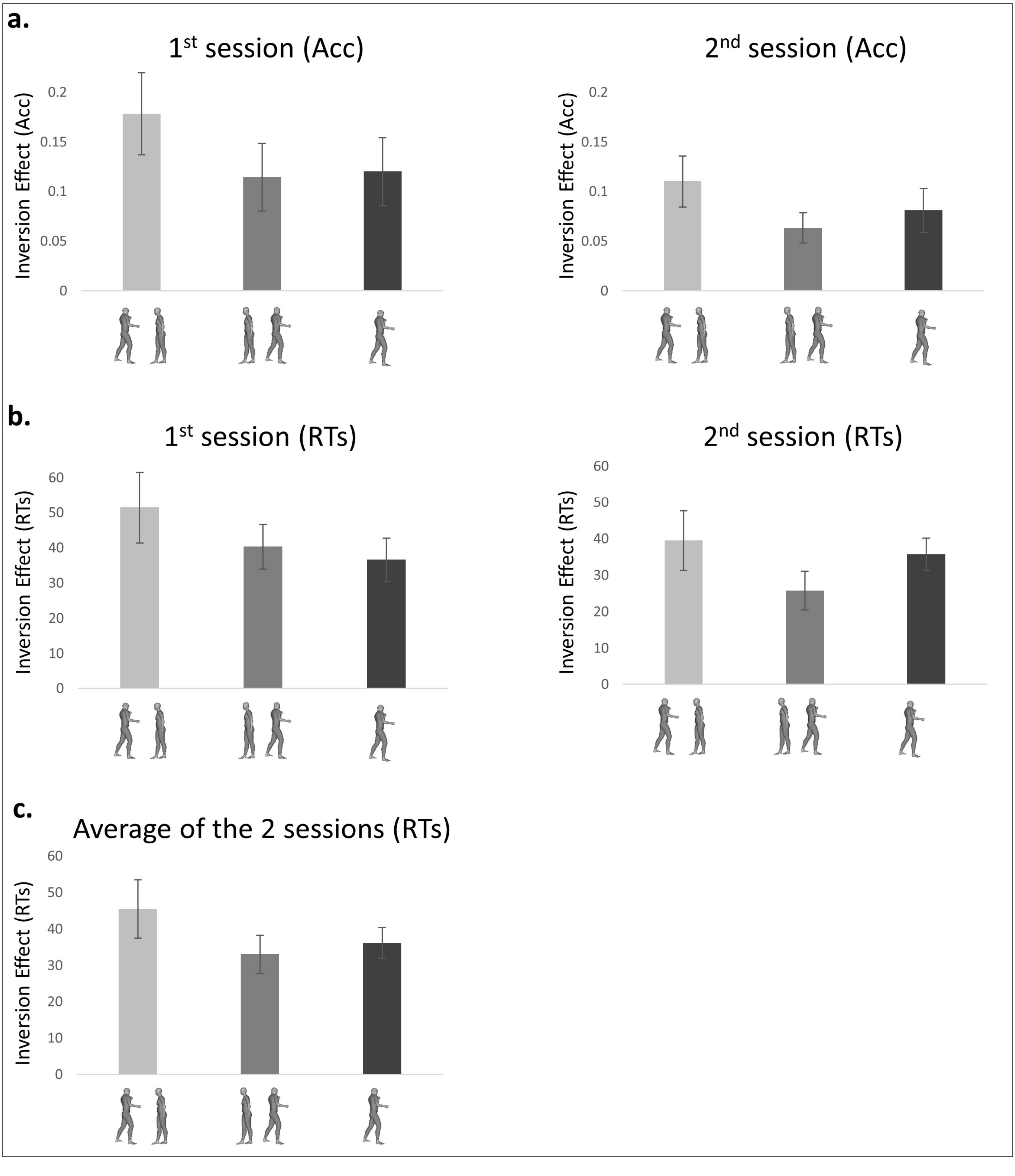

**Figure S1. Further accuracy and RTs results of the two session of the visual categorization task. a.** Inversion effect in the accuracy (proportion of correct response for upright minus inverted stimuli ± within-subjects normalized SEM) in the visual categorization of facing dyads, non-facing dyads and single bodies, in session 1 (left) and 2 (right) **b.** Inversion effect in the RTs (RTs for inverted minus upright stimuli ± within-subjects normalized SEM) in the visual categorization of facing dyads, non-facing dyads and single bodies, in session 1 (left) and 2 (right). **c.** Inversion effect in the RTs (± within-subjects normalized SEM) averaged over the two sessions.

**5. 2BIE brain-behavior correlation (TFCE corrected).** Following the whole-brain correlation between voxels responding to bodies and the 2BIE acquired during the first behavioral session, the statistical map of Pearson's correlation coefficients (Fisher-transformed) was corrected for multiple comparisons. To do so, we used a permutation testing approach (Nichols and Holmes, 2002) combined with threshold free cluster enhancement transformations (TFCE, Smith and Nichols, 2009) through the MatlabTFCE toolbox (permutation number = 5000, H = 2, E = 0.5, dh=0.1, Connectivity = 26, one-tailed). After correcting for multiple comparisons, a single cluster survived in the right occipital gyrus (peak MNI coordinates: 44 -82 -6; peak $z$-value = 2.09; peak $p$-value = 0.018; cluster size = 29).

**6. Correlation between the behavioral and neural 2BIE based on the behavioral session 1 only for left and right ROIs separated.** Using the measure of the 2BIE obtained in session 1 of the behavioral task, we ran Pearson's correlation with the neural 2BIE measured with fMRI. We found a significant correlation in both the right and left EBA (right: $r(20) = 0.49$, $p = 0.021$; left: $r(20) = 0.45$, $p = 0.037$). We did not find any correlation for the other ROIs (right FBA: $r(20) = 0.14$, $p > 0.250$; left FBA: $r(20) = 0.13$, $p > 0.250$; right FFA: $r(20) = 0.180$, $p > 0.250$ ; left FFA: $r(20) = 0.28$, $p = 0.211$; right PPA: $r(20) = -0.19$, $p > 0.250$; left PPA: $r(20) = 0.35$, $p = 0.113$; right EVC: $r(20) = 0.01$, $p > 0.250$; left EVC: $r(20) = -0.01$, $p > 0.250$).

**7. Correlation between the behavioral and neural 2BIE based on the behavioral Session 2.** Using the measure of the 2BIE obtained in session 2 of the behavioral task, we ran Pearson's correlation with the neural 2BIE measured with fMRI. We found a significant correlation in the right EBA ($r(20) = 0.49$, $p = 0.021$) (Fig. S2a), but not for the left EBA ($r(20) = 0.06$, $p > 0.250$). We didn't find any correlation for the other ROIs (right FBA: $r(20) = -0.05$, $p > 0.250$; left FBA: $r(20) = -0.09$, $p > 0.250$; right FFA: $r(20) = 0.26$, $p = 0.250$ ; left FFA: $r(20) = 0.01$, $p > 0.250$; right PPA: $r(20) = -0.09$, $p > 0.250$; left PPA: $r(20) = 0.07$, $p > 0.250$; right EVC: $r(20) = -0.07$, $p > 0.250$; left EVC: $r(20) = 0.03$, $p > 0.250$).

We then tested the correlation of the neural 2BIE computed for each voxel responsive to bodies across the brain, with the 2BIE measured in the behavioral session 2 (Fig. S2b). This analysis revealed a right lateralized cluster in the occipital gyrus, overlapping with the effect found with the correlation between the neural BIE and the BIE measured in the behavioral session 1 (peak MNI coordinates = 44 -86 -6, cluster size = 99, peak $z$-value = 3.42, peak $p$-value < 0.001). However, this cluster did not survive the correction for multiple comparisons.

**a.**

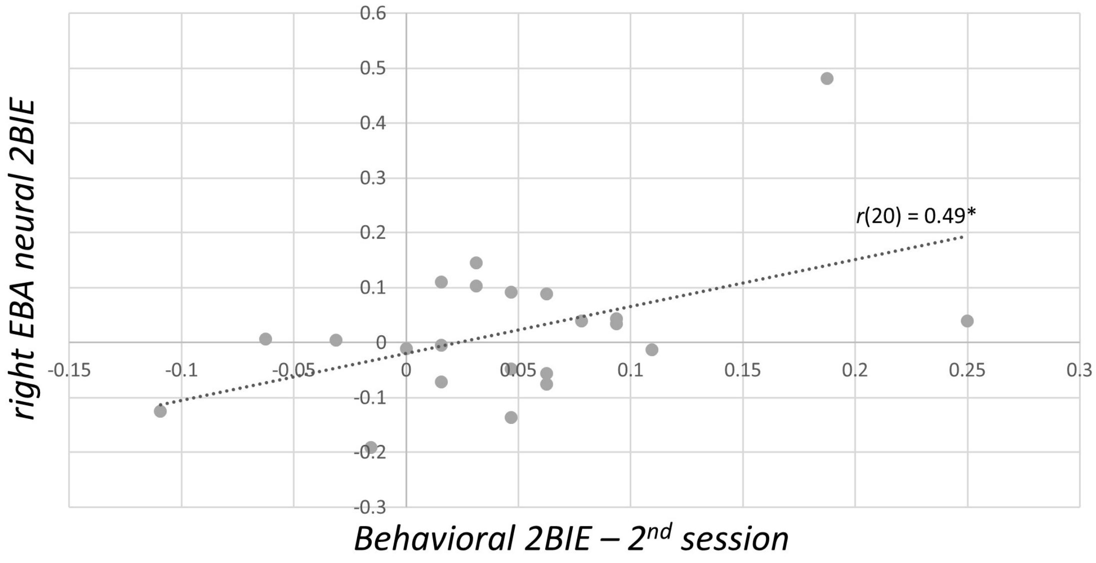


**b.**

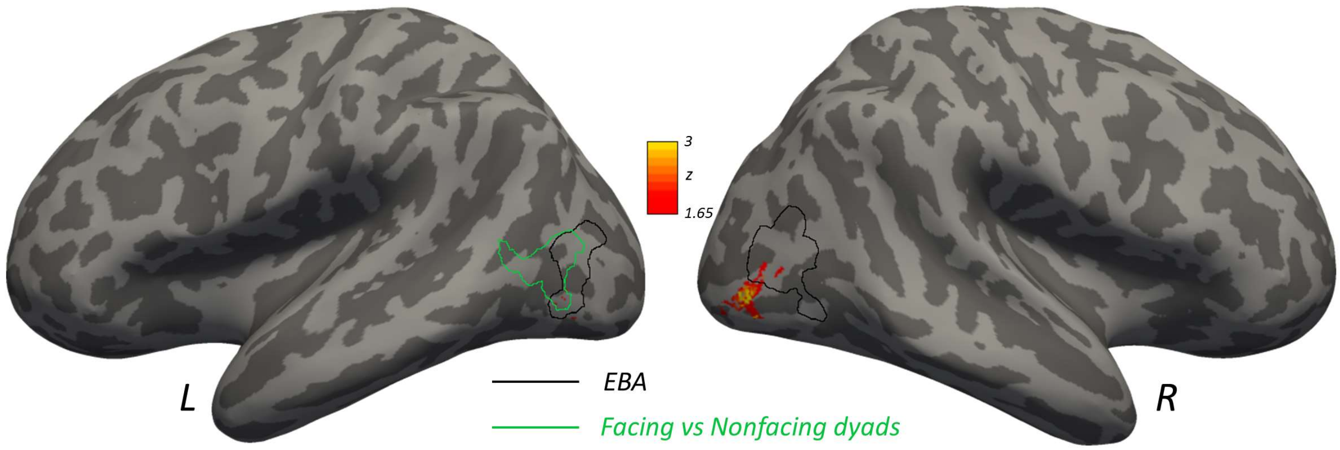


**Figure S2. Brain-behavior mapping for the 2BIE using data from session 2 of the behavioral task**. **a.** Correlation between the neural-2BIE within the functionally defined right EBA and the behavioral 2BIE measured in session 2 of the behavioral task. $*p \leq 0.05$ (one-tail). **b.** Statistical map representing the results of the correlation between the neural 2BIE and the behavioral 2BIE (session 2) computed in each body-responsive voxel across the whole brain. The color bar indicates $z$ values. The EBA location corresponds to the group-level random-effect contrast bodies > [objects+faces+places] using the data of the functional localizer task. The facing > nonfacing dyads cluster corresponds to the group-level random-effect contrast computed from data recorded during the main fMRI experiment.

### 8. Correlation between the behavioral and neural 2BIE based on the averaged performance over sessions 1 and 2.

Using data averaged across the two behavioral sessions for correlation with the neural BIE, we found a significant effect in the right EBA ($r(20) = 0.54$, $p = 0.009$), but not for the left EBA ($r(20) = 0.29$, $p = 0.183$). We did not find any correlation for the other ROIs (right FBA: $r(20) = 0.19$, $p > 0.250$; left FBA: $r(20) = 0.04$, $p > 0.250$; right FFA: $r(20) = 0.30$, $p = 0.181$ ; left FFA: $r(20) = 0.11$, $p > 0.250$; right PPA: $r(20) = -0.25$, $p > 0.250$; left PPA: $r(20) = -0.08$, $p > 0.250$; right EVC: $r(20) = -0.03$, $p > 250$; left EVC: $r(20) = 0.02$, $p > 0.250$).

We also tested the correlation of the neural 2BIE computed for each voxel responsive to bodies across the brain, with the 2BIE measured using the performance average of the session 1 and 2. This analysis revealed a right lateralized cluster in the occipital gyrus, overlapping with the effect found with the correlation between the neural BIE and the BIE measured in the behavioral sessions 1 and 2 (peak MNI coordinates = 44 -86 -6, cluster size = 7, peak $z$-value = 3.72, peak $p$-value < 0.001). However, this cluster did not survive the correction for multiple comparisons.

**9. Correlation between the 2BIE and AQ correlation based on the behavioral session 2 only and on the averaged performance over sessions 1 and 2.** There was a significant correlation between the AQ and the 2BIE when using the average performance across the two sessions of the behavioral task (Spearman $Rho(20) = -0.49$; $p = 0.021$), but not when only considering the subjects' performance in session 2 (Spearman $Rho(20) = -0.35$; $p = 0.101$).

### 10. Supplementary tables

**Table S1. Location and significance of clusters showing stronger responses to upright (UPR) vs. inverted (INV) orientation for facing and nonfacing dyads.**

| Contrast: UPR vs. INV | Hemispheres | Peaks locations | Peaks coordinates | | | *z* | Cluster-Wise FWE | Cluster size |
|---|---|---|---|---|---|---|---|---|
| | | | x | y | z | | | |
| Facing Dyads | Left | Early visual cortex | -10 | -100 | 12 | 6.21 | <0.001 | 1321 |
| | | Fusiform Gyrus | -40 | -40 | -20 | 5.01 | 0.003 | 302 |
| | | Lateral occipital cortex | -46 | -72 | 10 | 4.78 | <0.001 | 684 |
| | Right | Fusiform Gyrus | 42 | -42 | -22 | 5.67 | <0.001 | 461 |
| | | Lateral occipital cortex | 48 | -68 | 4 | 4.93 | <0.001 | 1129 |
| Nonfacing dyads | Left | Early visual cortex | -8 | -104 | 4 | 5.83 | <0.001 | 703 |
| | Right | Early visual cortex | 10 | -94 | 10 | 6.14 | <0.001 | 816 |
| | | Lateral occipital cortex | 42 | -58 | 14 | 4.46 | 0.002 | 456 |

**Table S2. Studies comparing inversion effects for facing dyads with inversion effects for faces, bodies, objects and nonfacing dyads** (data are presented visually in Figure 1 in the main text). This table is adapted from, and based on a meta-analysis in Rezlescu et al. (2016), to which we added data for facing and non-facing body-dyads, as measured in three previous studies and in the current one (red entries). Studies are sorted according to the scaled inversion effects found for faces, from smallest to largest, and for facing dyads, from smallest to largest. Scaled inversion effect = (inversion effect) / (1-chance).

| References | DATA | | | | Inversion Effect (% Acc) | | | | | Scaled Inversion (% Acc) | | | | | Object type |
|---|---|---|---|---|---|---|---|---|---|---|---|---|---|---|---|
| | Expt | Cond | n | Chance(%) | Faces | Bodies | Facing bodies | NonFacing bodies | Objects | Faces | Bodies | Facing bodies | NonFacing bodies | Objects | |
| Lahaie2006 | 1 | 1 | 16 | 50 | -1.1 | | | | 0.7 | -2.2 | | | | 1.4 | Greebles |
| Leder2006 | 1 | 1 | 36 | 16.7 | 1.8 | | | | 7.4 | 2.2 | | | | 8.9 | House |
| Leder2006 | 1 | 2 | 36 | 16.7 | 1.9 | | | | 1.8 | 2.3 | | | | 2.2 | House |
| Yin1969 | 3 | 1 | 23 | 50 | 1.3 | | | | 0.3 | 2.7 | | | | 0.6 | Costumes |
| Yarmey1971 | 1 | 1 | 80 | 50 | 2.2 | | | | 1.4 | 4.4 | | | | 2.8 | Dogs |
| | | | | | | | | | 0.9 | | | | | 1.8 | Buildings |
| deGelder2009 | 1 | 1 | 75 | 50 | 2.3 | | | | -0.6 | 4.6 | | | | -1.2 | Shoes |
| Haxby1999 | 1 | 1 | 6 | 50 | 2.8 | | | | 1.1 | 5.6 | | | | 2.2 | House |
| deGelder2015 | 1 | 1 | 32 | 50 | 3 | | | | -2 | 6 | | | | -4 | Shoes |
| Yin1970 | 1 | 1 | 12 | 50 | 3.3 | | | | 1.2 | 6.7 | | | | 2.3 | House |
| Yin1969 | 1 | 1 | 26 | 50 | 3.5 | | | | 1.2 | 6.9 | | | | 2.4 | House |
| | | | | | | | | | 0.2 | | | | | 0.4 | Airplane |
| Leder2006 | 1 | 6 | 36 | 16.7 | 6.5 | | | | 2.8 | 7.8 | | | | 3.4 | House |
| Boutet2006 | 2 | 2 | 15 | 50 | 4 | | | | 4 | 8 | | | | 8 | Chair/House |
| deGelder2015 | 1 | 2 | 26 | 50 | 4 | | | | -2 | 8 | | | | -4 | Shoes |
| Reed2006 | 3 | 1 | 24 | 50 | 4.9 | 2.3 | | | | 9.8 | 4.6 | | | | Bodies |
| Urgesi2014 | 1 | 1 | 12 | 50 | 5.7 | 3.7 | | | 3.7 | 11.3 | 21.6 | | | 7.3 | Motorcycles |
| Meinhardt-Injac2014 | 1 | 2 | 44 | 50 | 6 | | | | 0 | 12 | | | | 0 | Watch |
| Picozzi2009 | 2 | 1 | 21 | 50 | 6.5 | | | | 9.2 | 13.1 | | | | 18.4 | Cars |
| Rossion2002 | 1 | 2 | 10 | 50 | 7 | | | | -2 | 14 | | | | -4 | Greebles |
| Reed2003 | 2 | 1 | 18 | 50 | 7 | 0 | | | | 14 | 10 | | | | Bodies |
| Boutet2006 | 2 | 1 | 15 | 50 | 7 | | | | 0 | 14 | | | | 0 | Chair/House |
| Yovel2008 | 1 | 1 | 74 | 50 | 8 | | | | 0 | 16 | | | | 0 | House |
| Abassi2020 | 1 | | | 50 | | 5.5 | 8 | 4 | 1.3 | | 11 | 16 | 8 | 2.6 | Chairs |
| Yovel2004 | 2 | 1 | 74 | 50 | 9 | | | | -4 | 18 | | | | -8 | House |
| Robbins2007 | 3 | 2 | 20 | 50 | 10 | | | | 2 | 20 | | | | 4 | Dogs |
| Leder2006 | 1 | 5 | 36 | 16.7 | 17.6 | | | | 3.7 | 21.1 | | | | 4.4 | House |
| Bosbach2006 | 1 | 1 | 12 | 50 | 11.1 | 2.8 | | | 2.8 | 22.2 | 17 | | | 5.6 | House |
| Leder2006 | 1 | 4 | 36 | 16.7 | 18.5 | | | | 4.6 | 22.2 | | | | 5.5 | House |
| Susilo2013 | 1 | 1 | 20 | 50 | 11.2 | 0 | | | | 22.4 | 19.2 | | | | Bodies |
| Brandman2012 | 1 | 1 | 98 | 50 | 12 | 0 | | | | 24 | 24 | | | | Bodies |
| Picozzi2009 | 1 | 1 | 36 | 50 | 12.4 | | | | -2.2 | 24.8 | | | | -4.4 | Shoes |
| Picozzi2009 | 3 | 1 | 10 | 50 | 13 | | | | -2.6 | 26 | | | | -5.2 | Cars |
| Picozzi2009 | 3 | 2 | 10 | 50 | 13 | | | | 2 | 26 | | | | 4 | Cars |
| Carey1977 | 1 | 1 | 36 | 50 | 13.3 | | | | 6.3 | 26.7 | | | | 12.7 | House |
| Yovel2010 | 1 | 1 | 10 | 50 | 14 | 0 | | | | 28 | 28 | | | | Bodies |
| Bushmakin2014 | 2 | 1 | 60 | 25 | 21 | | | | 14 | 28 | | | | 18.7 | Novel |
| Meinhardt-Injac2014 | 1 | 1 | 44 | 50 | 14 | | | | 0 | 28 | | | | 0 | Watch |
| Picozzi2009 | 2 | 2 | 19 | 50 | 14.7 | | | | -0.5 | 29.4 | | | | -0.9 | Cars |
| Valentine1986 | 3 | 2 | 16 | 50 | 14.8 | | | | 6.4 | 29.6 | | | | 12.8 | House |
| Bruce1991 | 6 | 1 | 62 | 50 | 15 | | | | 10 | 30 | | | | 20 | House |
| Yovel2004 | 2 | 2 | 74 | 50 | 15 | | | | -1 | 30 | | | | -2 | House |
| Yovel2008 | 1 | 2 | 74 | 50 | 15 | | | | -2 | 30 | | | | -4 | House |
| Papeo2019 | 3 | | 81 | 50 | | | 15.3 | 10.3 | 3 | | | 30.6 | 20.6 | 6 | Chairs |
| Papeo2017 | 3 | | 62 | 50 | | | 15.7 | 11.3 | 7 | | | 31.4 | 22.6 | 14 | Chairs |
| Valentine1986 | 2 | 1 | 20 | 50 | 16.6 | | | | 5.4 | 33.2 | | | | 10.8 | House |
| Busigny2010 | 4 | 1 | 12 | 50 | 17 | | | | 1.1 | 34 | | | | 2.2 | Cars |
| Leder2006 | 1 | 3 | 36 | 16.7 | 29.6 | | | | 0.9 | 35.5 | | | | 1.1 | House |
| Current study | 1 | | 22 | 50 | | 12 | 17.8 | 11.4 | 2.1 | | 24 | 35.6 | 22.8 | 4.2 | Chairs |
| Rossion2010 | 1 | 2 | 20 | 50 | 18 | | | | 4 | 36 | | | | 8 | Cars |
| Diamond1986 | 1 | 1 | 16 | 50 | 19 | | | | 9 | 38 | | | | 18 | Scenes |
| Bruyer1992 | 1 | 2 | 16 | 50 | 19 | | | | 4 | 38 | | | | 8 | Handwriting |
| Boutet2006 | 1 | 2 | 14 | 50 | 19 | | | | 3 | 38 | | | | 6 | Chair/House |
| Williams2007 | 1 | 1 | 32 | 50 | 19.5 | | | | 6.3 | 39 | | | | 12.6 | Radio |
| Phillips1979 | 1 | 1 | 95 | 50 | 19.7 | | | | 7.8 | 39.4 | | | | 15.6 | Woodcuts |
| Diamond1986 | 2 | 2 | 16 | 50 | 20 | | | | 8 | 40 | | | | 16 | Dogs |
| Boutet2006 | 1 | 1 | 14 | 50 | 20 | | | | 4 | 40 | | | | 8 | Chair/House |
| Valentine1986 | 3 | 1 | 16 | 50 | 21.8 | | | | 4.8 | 43.6 | | | | 9.6 | House |
| Diamond1986 | 3 | 2 | 16 | 50 | 23 | | | | 2 | 46 | | | | 4 | Dogs |
| Robbins2007 | 1 | 2 | 15 | 50 | 24 | | | | 3 | 48 | | | | 6 | Dogs |
| Busigny2010 | 3 | 1 | 9 | 50 | 25.3 | | | | 1 | 50.6 | | | | 2 | Cars |
